# Genetic Polymorphisms and Molecular Mechanisms Mediating Oncolytic Potency of Reovirus Strains

**DOI:** 10.1101/569301

**Authors:** Adil Mohamed, Derek R. Clements, Prathyusha Konda, Shashi A. Gujar, Patrick W. Lee, James R. Smiley, Maya Shmulevitz

## Abstract

The Dearing strain of Mammalian orthoreovirus (T3D) is undergoing clinical trials as an oncolytic virotherapeutic agent. In this study, a comprehensive phenotypic and genetic comparison of T3D virus stocks from various laboratories and commercial sources revealed that T3D laboratory strains differ substantially in their oncolytic activities *in vitro* and *in vivo*. Superior replication of the most-oncolytic T3D lab strain was attributed to several mechanistic advantages: virus-cell binding, viral RNA transcriptase activity, viral inclusion morphology, and differential activation of RIG-I versus NFκB-dependent signalling pathways. Viral S4, M1 and L3 gene segments were each independently associated with a distinct mechanistic advantage. Furthermore, the specific missense polymorphisms that governed replication potency were identified, and utilized to generate a hybrid of T3D laboratory strains with further-augmented replication in tumor cells. Together, the results depict an elaborate balance between reovirus replication and host-cell signaling to achieve optimal oncolytic reovirus efficacy.

## INTRODUCTION

Cancer is a global health concern and the necessity for novel cancer therapies is well appreciated. Viruses that specifically target tumor cells offer distinct benefits over conventional drug-based therapies. First, their amenability to engineering permits addition of multiple layers of specificity and tumoricidal activities, leading to low toxicity at high doses (Russell and Peng, 2018). Second, since viruses inherently replicate, their doses increase in tumors unlike conventional therapies. Third is that in addition to killing tumor cells directly, viruses induce secretion of cytokines and recruitment of immune cells to tumors, leading to promotion of anti-tumor immunity. These features of viruses that make them good potential therapies for cancer also come with a caveat; the immune system can clear viruses. Therefore, it is essential for an oncolytic virus to be so efficient that it can exert tumoricidal and immune-stimulating effects before clearance.

Three oncolytic viruses currently progressing through human phase III clinical trials include reovirus (REOLYSIN^®^), vaccinia virus (Pexa-Vec^®^) and herpes simplex virus (T-VEC/IMLYGIC^®^) (Amgen Inc, 2018; Oncolytics Biotech Inc, 2018; SillaJen Inc, 2018). T-VEC (Imlygic™) received FDA approval for “use in melanoma with injectable but non-resectable lesions in the skin and lymph nodes” (Pol et al., 2016). HSV and vaccinia are both large DNA viruses, and their tumor specificity was conferred by deleting viral genes, for example genes critical for replication that can be uniquely supplied by tumor cells but not normal cells, and/or genes that normally regulate innate and adaptive immune responses (Potts et al., 2017). In contrast, reovirus is a smaller segmented double-stranded RNA (dsRNA) virus that naturally replicates preferentially in tumor cells (Coffey et al., 1998; Hirasawa et al., 2002; Norman et al., 2002b; Wilcox et al., 2001). In other words, the unmodified reovirus promotes tumor clearance without causing toxicity to healthy tissues. Previous studies demonstrated that replication of reovirus in non-transformed cells is restricted through 4 mechanisms: (1) poor uncoating of outer capsid proteins, (2) production of progeny virions with reduced specific infectivity, (3) reduced cell death and virus release, and (4) high interferon antiviral responses (Marcato et al., 2007; Shmulevitz et al., 2005, 2010a; Shmulevitz et al., 2010b). The existence of multiple barriers in non-transformed cells ensures that reovirus replication is tumor-specific.

Although generally referred to as ‘reovirus’, the specific reovirus undergoing clinical trials is the serotype 3 Dearing isolate of mammalian orthoreoviruses (T3D, tradename Reolysin®). In Phase I-III clinical trials, reovirus T3D is well tolerated and demonstrates some indications of efficacy (Carlson et al., 2005; Oncolytics Biotech Inc, 2018; Stoeckel and Hay, 2006). But despite promise, reovirus therapy produces incomplete response in the majority of immunocompetent tumor-bearing animals (Gujar et al., 2014; Rajani et al., 2016; Shmulevitz et al., 2012) and human patients. To enhance the oncolytic activity of reovirus, researchers and pharmaceutical companies are testing combination therapy, for example with chemotherapies or checkpoint blockade inhibitors. To complement these efforts, our laboratory strives to understand the impact of virus features, such as replication proficiency in tumors, on therapeutic efficacy. We predicted that maximum replication of reovirus in tumors would promote both direct killing and immunotherapeutic activities. Moreover, we predicted that features of the virus will impact the immune-stimulatory landscape (e.g. cytokines), by interacting with specific cellular signaling pathways. To test these predictions, we explored the relationships between oncolytic activity and virus genetic and phenotypic features, among various highly related laboratory strains of T3D.

In this study, we discovered that highly related laboratory strains of T3D exhibit significant differences in oncolytic potency *in vitro* and *in vivo*. These findings reinforce that virus genomic and phenotypic features have a strong impact on oncolytic proficiencies. Since the T3D strains were highly genetically related, they provided an opportunity to discover novel determinants of reovirus oncolysis. A comprehensive genetic and phenotypic comparison of T3D laboratory strains revealed that the most-oncolytic strain (obtained from the Patrick Lee laboratory, T3D^PL^) has 3 distinct and additive advantages over the less-oncolytic strains (obtained from the Terence Dermody and Kevin Coombs laboratories, T3D^TD^ and T3D^KC^, respectively). Relative to T3D^TD^ and T3D^KC^, T3D^PL^ exhibited superior replication in transformed cells in a single round of infection, and induced less IFN signaling which facilitated enhanced cell-cell spread. The genes that contribute to enhanced replication and spread of T3D^PL^ were then characterized, and found to be (1) the T3D^PL^ S1-encoded σ1, which increased the proportion of virions that harbour sufficient number of σ1 trimers for binding to cells, (2) T3D^PL^ S4-encoded σ3, which increased expression of NF-κB stimulated genes, (3) the M1-encoded μ2, which increased virus transcription and filamentous factories, and (4) the L3-encoded λ1, which increased virus factory size. When evaluating which specific polymorphisms in S4, M1, and L3 contribute to oncolysis, surprisingly specific amino acid differences between T3D^PL^ versus T3D^TD^ had both positive and negative contributions, and ultimately a hybrid between T3D^PL^ and T3D^TD^ exhibited superior oncolytic activity even relative to T3D^PL^. This finding suggests that the T3D oncolytic vector may benefit from continued genetic development to fully thrive in tumors. Finally, important to the current excitement surrounding the immunotherapeutic value of oncolytic viruses, was our discovery that T3D laboratory strains induced expression of different cytokines by infected tumor cells. Specifically, while T3D^PL^ caused high induction of NF-κB-dependent cytokines but low induction of IFN-dependent cytokines, the reciprocal was true for T3D^TD^. Altogether the study reveals novel strategies of improving specific steps of virus replication, as well as cytokine response, to augment oncolytic potency.

## RESULTS

### Laboratory strains of reovirus exhibit distinct oncolytic activity *in vitro* and *in vivo*

As described above, REOLYSIN^®^ is a promising oncolytic agent that is undergoing numerous clinical trials. REOLYSIN^®^ was derived from a stock of T3D obtained from the laboratory of Patrick Lee (T3D^PL^). T3D strains from different laboratories were previously shown to differ in several properties including factory morphology and *in vivo* respiratory pathogenesis, but T3D^PL^ (or Reolysin^®^) were not included in these studies, nor was replication efficiency in transformed cells suggested to be different (Nygaard et al., 2013; Parker et al., 2002; Yin et al., 2004). The use of distinct laboratory origins by laboratories across the world for studies on reovirus oncolysis made it imperative to determine if all strains of T3D share similar oncolytic properties. Accordingly, we compared the *in vitro* and *in vivo* oncolytic properties of T3D^PL^ to those of T3D obtained from Kevin Coombs (T3D^KC^), Terence Dermody (T3D^TD^), and the American Type Culture Collection (T3D^ATCC^).

*In vitro* oncolysis by a virus is defined by its ability to infect, disseminate, and ultimately cause ‘destruction or dissolution’ (oncolysis) of tumorigenic cells. As an initial strategy to explore *in vitro* oncolysis, we compared the plaque size of the various T3D laboratory strains on L929 tumorigenic mouse fibroblasts. Plaque size provides a measure of overall efficiency of virus replication and cell-to-cell spread over several rounds of infection. CsCl-purified virus preparations were subjected to plaque titration and visualized by crystal violet staining (Figure 1A). Plaque size was quantified by densitometry and reported as units of pixels. When compared at 7 days post-infection (dpi), T3D^PL^ had the largest average plaque size of 1644 pixels. T3D^KC^ and T3D^TD^ had the smallest average plaque sizes of 42 pixels and 32 pixels, respectively. T3D^ATCC^ had an intermediate average plaque size of 225 pixels. To our knowledge, this is the first report of differential virus replication and *in vitro* oncolysis among T3D laboratory strains.

**Figure 1.**
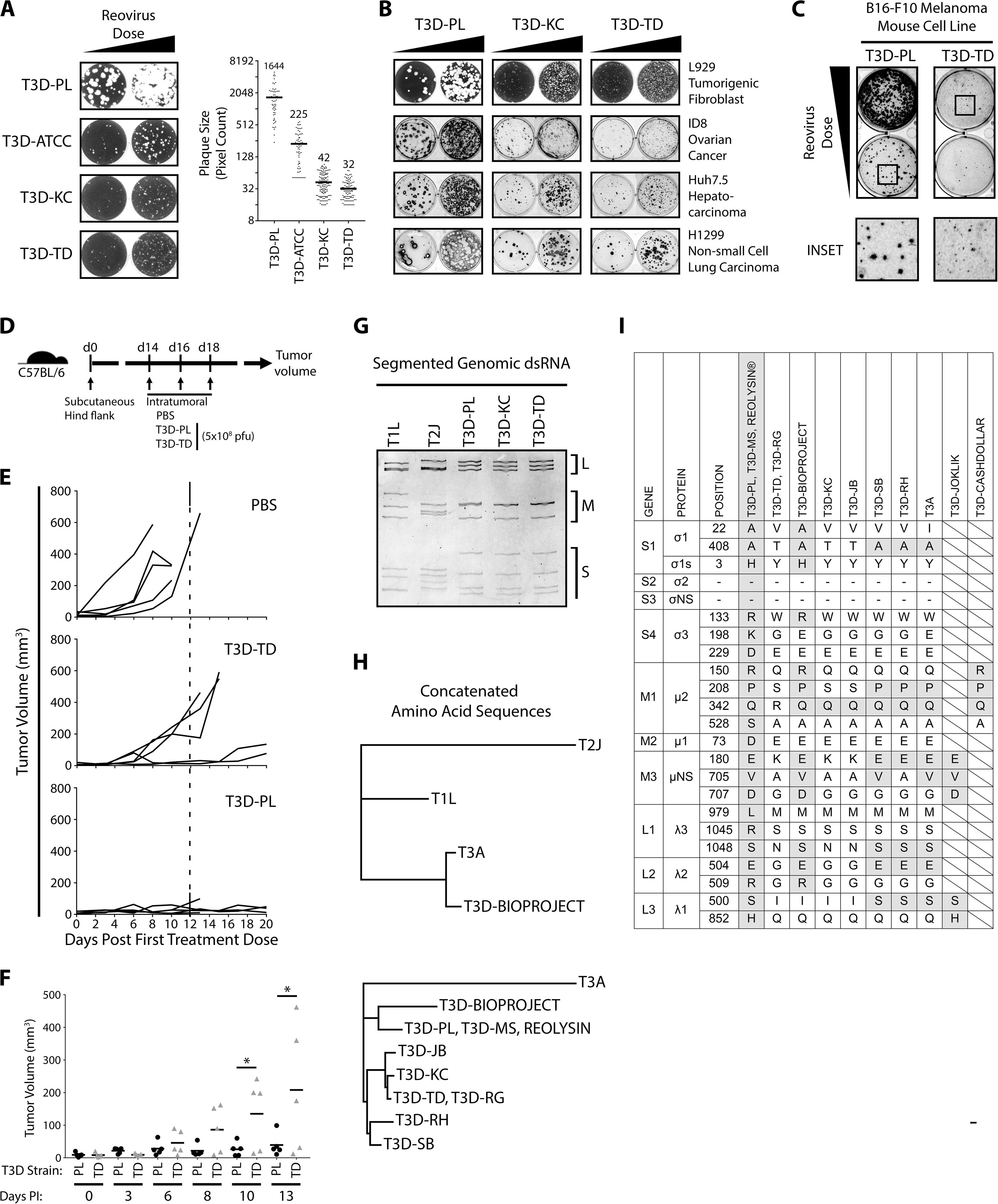
T3D^PL^ Laboratory Clone has Enhanced Oncolytic Activity *In Vitro* and *In Vivo*. **(A)** Plaques generated by reovirus T3D strains (T3D^PL^, T3D^ATCC^, T3D^KC^ and T3D^TD^) on L929 cell line. Reovirus plaques were visualized by crystal violet staining (left) and plaque size quantified by ImageQuant colony counting software (right). Mean plaque size indicated above data points. **(B)** Plaques and foci generated by reovirus T3D strains (T3D^PL^, T3D^KC^ and T3D^TD^) on L929, ID8, Huh7.5 and H1299 cell lines. Reovirus plaques (L929) were visualized with crystal violet and reovirus foci (ID8, Huh7.5 and H1299) were stained with colorimetric immunocytochemistry using primary polyclonal reovirus antibody, alkaline phosphatase secondary antibody and BCIP/NBT substrate. **(C)** T3D^PL^ and T3D^TD^ foci formation on B16-F10 cells visualized by colorimetric immunocytochemistry as in (B). Inset displays a magnification of indicated well area. **(D)** Schematic timeline for comparing oncolytic potency of reovirus T3D strains in the B16-F10 mouse melanoma model. **(E)** Following the first intratumoral inoculation of PBS, T3D^PL^ or T3D^TD^, tumor volume was monitored using caliper measurements every 2-3 days. n=5 mice per group. **(F)** Tumor volume comparison from 0-13 days post initial T3D^PL^ or T3D^TD^ dose. Data points obtained from (E). Statistical significance determined using unpaired t test, * p<0.05. **(G)** Separation of reovirus dsRNA segments using SDS-PAGE and RNA staining using ethidium bromide. Genomic reovirus dsRNA was Trizol LS extracted from CsCl purified reovirus preparations. **(H)** Reovirus phylogenetic trees generated with concatenated complete reovirus amino acid sequences using the Geneious^®^ tree builder add on. Assembly of phylogenetic trees was performed using the Jukes-Cantor genetic distance model and neighbour-joining tree build method. Outgroup set as T2J (top) or T3A (bottom). Concatenate sequences of 11 reovirus proteins, in order σ1, σ1s, σ2, σNS, σ3, μ1, μ2, μNS, λ1, λ2, λ3. **(I)** Specific amino acid differences between T3DPL and T3DTD. Amino acids at respective positions for T3D^BIOPROJECT^, T3D^KC^, T3D^JB^, T3D^SB^, T3D^RH^, T3A, T3D^JOKLIK^, T3D^CASHDOLLAR^ are included for source laboratory divergence.

T3D^PL^ can be traced back to the Joklik, W.K. laboratory (Duke University). Both T3D^KC^ and T3D^TD^ laboratory strains originated from the Fields, B.N. laboratory (Harvard Medical School), which was initially obtained from the Joklik laboratory but was proposed to have diverged from other T3D strains by the mid-1970s (Cross and Fields, 1972; Fields, 1971). The depositor of T3D^ATCC^, W. Adrian Chappell, was affiliated with the Centre for Disease Control (Atlanta), but could not be linked to any reovirus laboratories or publications. Due to the uncertain origin of T3D^ATCC^, we omitted this strain from our subsequent experiments. However, laboratories using T3D^ATCC^ should note that replication kinetics are distinct from those of other T3D laboratory strains.

To determine if differential *in vitro* oncolysis by T3D strains extend to other cancer cells, we compared plaque sizes on a panel of mouse (ID8 ovarian cancer and B16-F10 melanoma) and human (Huh7.5 hepatocellular carcinoma and H1299 non-small cell lung carcinoma) tumorigenic cell lines (Figure 1B and 1C). Immunocytochemical staining with polyclonal anti-reovirus antibodies was used to monitor reovirus-infected plaques, as not all cells produce monolayers sufficiently dense for crystal violet staining. In all cell lines assessed, T3D^PL^ formed visibly larger plaques compared to T3D^KC^ and T3D^TD^. Additionally, clearing of cells in the center of reovirus-infected plaques, indicative of cell death, was more apparent for T3D^PL^ relative to T3D^KC^ and T3D^TD^. Therefore, relative to T3D^KC^ and T3D^TD^, T3D^PL^ displayed greater *in vitro* oncolytic potency on an assortment of cancer cells.

Few studies have directly assessed whether *in vitro* oncolysis by viruses correlates with *in vivo* oncolytic activity, and hence it was critical to determine whether differences in *in vitro* replication potency corresponded with differential oncolytic efficacies *in-vivo*. T3D^PL^ was previously characterized to reduce tumor burden in the aggressive B16-F10 syngeneic melanoma mouse model (Shmulevitz et al., 2012) and therefore this model was chosen to compare the *in-vivo* oncolytic efficacy of T3D^PL^ versus T3D^TD^. B16-F10 cells were implanted subcutaneously into the hind flank of six-week old female C57BL/6 mice (Figure 1D). When tumors were palpable, equivalent doses (5×10^8^ pfu/dose) of T3D^PL^ and T3D^TD^ were injected intra-tumorally at 2-day intervals, for a total of 3 doses. PBS was injected as a negative control. Tumor volumes were monitored every 2 days post-inoculation. Compared to the PBS control group, both T3D^PL^ and T3D^TD^ caused delayed tumor growth (Figure 1E). However, T3D^TD^ only marginally controlled tumor growth while T3D^PL^ controlled tumor growth in all mice. Specifically, at 13 days post inoculation when all mice were alive, T3D^PL^-treated animals had significantly smaller tumors (P=0.0024, n=5) than T3D^TD^-treated mice (Figure 1F). Animal survival could not be compared because, as per the strict animal ethics protocol, mice had to be sacrificed when tumors became too large (~600mm^3^), or showed signs of necrosis or ulceration. While 3/5 T3D^TD^-treated mice were sacrificed due to large tumor volumes, two T3D^PL^-treated mice were sacrificed due to tumor scabbing despite small tumor volume and likely a positive indication of oncolytic activity. Thus, survival curves were uninformative with respect to reovirus mediated tumor regression. Importantly, no signs of virus pathogenesis in normal tissues were observed; specifically, reovirus can cause black tail and ears, cardiotoxicity and neurotoxicity in immunodeficient mice due to replication at sites other than tumors (Kim et al., 2011; Loken et al., 2004), but no such symptoms were observed in our immunocompetent mice. These results demonstrate that for reovirus, robust virus infection *in vitro* correlates with efficient *in vivo* oncolysis. These findings are important because a recent focus on the immunostimulatory aspect of oncolytic viruses has put into question the importance of virus replication proficiency in tumors as a determinant of oncolytic potency; our results demonstrating a strong relationship between virus replication *in vitro* and oncolysis *in vivo* for reovirus.

### Genetic variations between the differentially-oncolytic T3D laboratory strains

The differences in oncolytic potency among T3D lab strains warranted a close analysis of genetic divergence between T3D stocks passaged in independent laboratories. First, resolution of reovirus genomic dsRNA segments using SDS-PAGE was used to determine if gross changes could be observed among the 10 reovirus genome segments. Reovirus serotypes 1, 2, and 3 produce unique electrophenotype (Coombs, 2011; Shatkin et al., 1968), but even independent isolates of reovirus from the same serotype commonly show mobility differences (Hrdy et al., 1979). T3D^PL^, T3D^KC^ and T3D^TD^ electrophenotypes were indistinguishable, although very distinct from serotype 1 Lang (T1L) and serotype 2 Jones (T2J) prototypes (Figure 1G), suggesting close ancestry.

Extensive genetic and phenotypic characterization of reovirus has been performed by comparing prototypic reovirus serotypes, commonly T1L versus T3D, but an in-depth analysis of whole genome sequences of T3D laboratory strains has yet to be performed. We collected and assembled complete reovirus nucleotide and amino acid sequences available on the NCBI database. T3D^BioProject^ represents the earliest published complete T3D sequences, which were obtained as a collaborative initiative as per NCBI BioProject PRJNA14861. As mentioned, REOLYSIN® is the proprietary reovirus strain under clinical investigation for cancer therapy, with sequence data published by Oncolytics Biotech (Chakrabarty et al., 2014). T3D^JB^ (John Bell laboratory), T3D^RH^ (Rob Hoeben laboratory), serotype 3 Abney (T3A), T1L, and T2J sequences from NCBI database were also included. The complete sequences for T3D^TD^ were obtained from Terence Dermody but also confirmed in-house to ensure absence of divergence in our hands. Finally, our laboratory strain T3D^MS^ obtained from T3D^PL^ over 15 years ago, T3D^TD^, T3D^PL^ and T3D^KC^ were also sequenced in-house.

For an overall reovirus lineage assessment, we used concatenated sequences of 11 reovirus proteins (σ1, σ1s, σ2, σNS, σ3, μ2, μ1, μNS, λ3, λ2, λ1) from each reovirus strain to generate a phylogenetic tree using the Jukes-Cantor model and Neighbour-Joining method (Geneious®). All T3D strains branched together independent of the other reovirus serotypes T1L, T2J and T3A (Figure 1H, top). T3D^PL^ and REOLYSIN® had 6 nucleotide differences but identical amino acid sequences, confirming homogeneity of the laboratory strain originating from Patrick Lee. T3D^PL^ and REOLYSIN® branched separately from T3D^TD^ and T3D^KC^ (Figure 1H, bottom). Other laboratory strain sequences such as T3D^JB^, T3D^RH^, and T3D^SB^ clustered more-closely to T3D^TD^ and T3D^KC^ than to T3D^PL^.

To define the specific differences among T3D laboratory strains, we aligned the individual protein sequences (Figure 1I). T3D^PL^ and REOLYSIN® were the most similar strains to T3D^BioProject^ in σ1, σ1s, μ2, μNS and λ2 proteins. T3D^TD^ and T3D^KC^ were most similar to T3D^BioProject^ in μ1, λ1 and λ3 proteins. All T3D strains had identical σ2 and σNS proteins. Interestingly, the BioProject collaboration had σ1, σ1s, μ2, μNS and λ2 sequences obtained from the Wolfgang Joklik-affiliated laboratories, while μ1, λ1 and λ3 protein sequences were obtained from the Bernard Fields-affiliated laboratories. Accordingly, we predict the major distinction between laboratory strains of T3D dates as far back as the conception of the Bioproject. Altogether there were 43 and 45 nucleotide differences between T3D^PL^ versus T3D^TD^ or T3D^KC^, respectively, of which 21 and 22 resulted in amino acid changes. The amino acid differences were dispersed among 8 of the 10 virus genome segments.

### The S4, M1 and L3 genes are determinants of enhanced T3D^PL^ oncolysis

The distinct oncolytic potencies of T3D^PL^ versus T3D^TD^ despite only 21 amino acid differences provided a unique opportunity to decipher the genetic and phenotypic basis for robust oncolysis. As the first step, a classical genome segment reassortment analysis was conducted in search of reovirus genes that contribute to the enhanced plaque size of T3D^PL^. L929 cells were co-infected with T3D^PL^ and T3D^TD^ at a ratio of 1:10 to favor reassortment of T3D^PL^ genes in a primarily T3D^TD^ background (Figure 2A). The co-infection lysates were plaque titrated and 20 large plaques were twice plaque purified and sequenced. Of the 20 large plaques, 8 were of composed entirely of T3D^PL^ genes (Figure 2B). The remaining 14 large plaques were reassortants that contain between 2-7 T3D^PL^ genome segments. The T3D^PL^-derived M1 gene was present in all (12/12) reassortants, and was therefore very likely a determinant of the larger plaque size of T3D^PL^. Other T3D^PL^ genome segments found in ≥50% of the large-plaque reassortants were S4 (9/12) and L3 (6/12).

**Figure 2.**
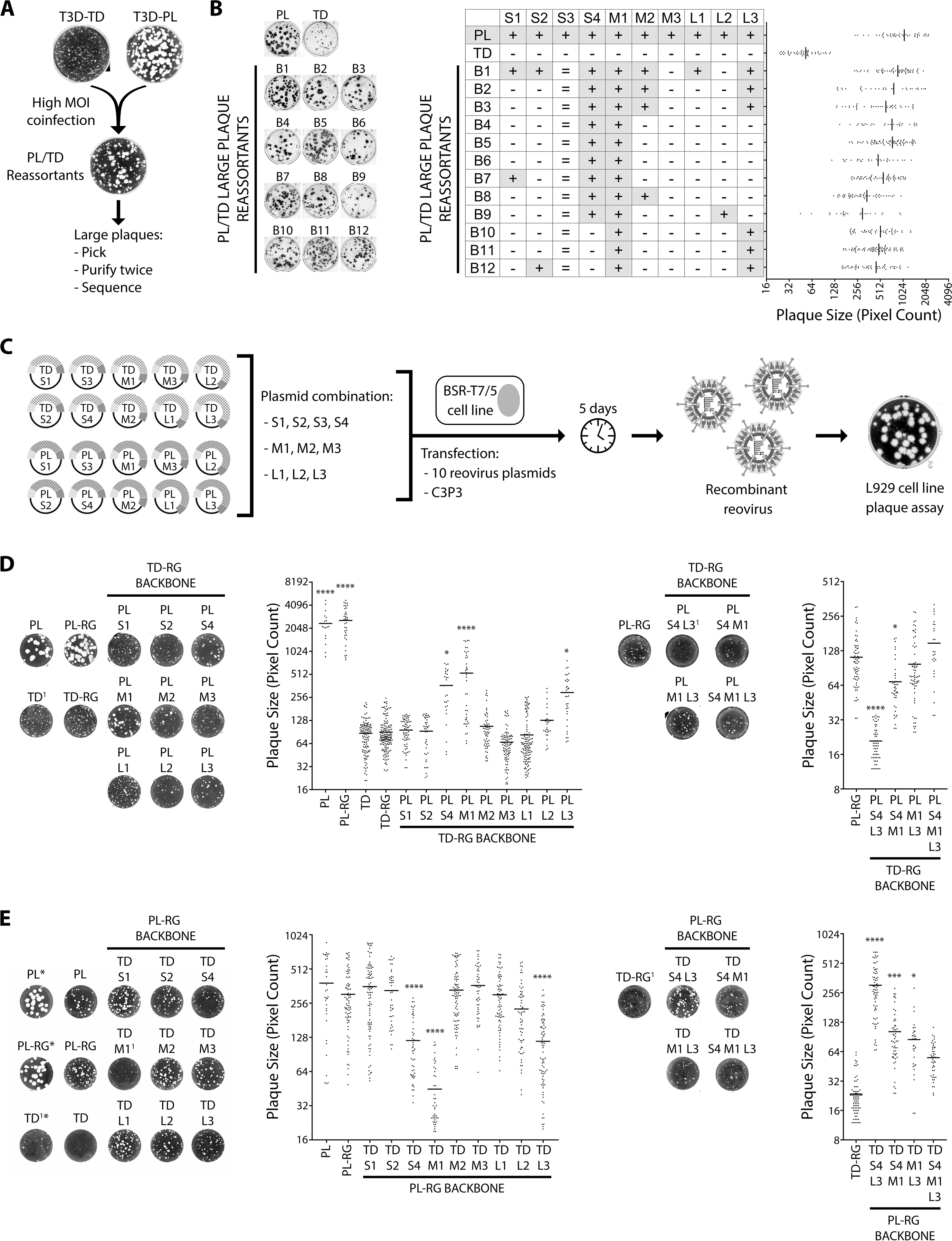
PL-S4, PL-M1 and PL-L3 genes contribute towards enhanced T3D^PL^ plaque size. **(A)** L929 cells were coinfected T3D^PL^ and T3D^TD^ at high MOI (T3D^PL^ MOI 20, T3D^TD^ MOI 200). At 24hpi, cell lysate was collected and following 3 freeze/thaw cycles, plaque assays were performed on L929 cells. Using parental T3DPL and T3DTD plaques as a reference, twenty plaques similar in size to T3DPL were isolated, twice plaque purified on L929 cells and sequenced. **(B)** (Left) Foci size comparison of parental (PL and TD) and large plaque reassortants (B1-B12) following colorimetric immunocytochemical staining using primary polyclonal reovirus antibody, alkaline phosphatase secondary antibody and BCIP/NBT substrate. (Right) Summary of sequencing analysis of T3D^PL^/T3D^TD^ reassortants indicating parental gene (+ T3D^PL^, – T3D^TD^) matched to plaque area quantification using ImageQuantTL colony counting add-on. The S3 gene was identical (=) between T3D^PL^ and T3D^TD^. **(C)** BHK/T7-9 cells were transfected with a combination of plasmids encoding10 reovirus gene segments and C3P3. At 5 days post transfection, cell lysates were harvested, and recombinant reovirus plaque size was determined on L929 cells. C3P3 plasmid encodes T7 RNA polymerase and African swine fever virus NP868R capping enzyme separated by a linker sequence. **(D)** and **(E)** Using L929 cells, plaques of parental and recombinant reoviruses were visualized following crystal violet staining and quantified using ImageQuantTL colony counting add-on. (D) TD backbone with PL gene combinations (E) PL backbone with TD gene combinations. PL and TD are purified laboratory stocks of T3D^PL^ and T3D^TD^, respectively. PL-RG and TD-RG are reverse genetics generated virus lysates using complete set of parental T3D^PL^ and T3D^TD^ plasmids, respectively. Statistical significance determined using one-way ANOVA with Dunnett's multiple comparisons test, * p < 0.05, *** p < 0.001, **** p < 0.0001. Samples in (D) compared to TD-RG. Samples in (E) compared to PL-RG. ^1^ Set of plaques were stopped when plaques in this well were visible

To further discriminate genes that contribute to the large plaque phenotype of T3D^PL^, we generated mono-reassortants between T3D^PL^ and T3D^TD^ using reverse genetics (RG) (Figure 2C). The 10 reovirus genome segments from T3D^PL^ and T3D^TD^ were cloned in-house into the T7 RNA polymerase-driven reverse genetics plasmid backbone. Parental viruses generated using the RG system (PL^RG^ and TD^RG^) produced plaques of similar size to T3D^PL^ and T3D^TD^, respectively, confirming genetic authenticity of reverse genetics-derived viruses (Figure 2D and 2E). Mono-reassortants in a TD^RG^ backbone revealed that T3D^PL^-derived M1, S4 and L3 significantly increased plaque size compared to TD^RG^ (Figure 2D). Importantly, none of the PL genes individually restored plaque size to the parental PL^RG^. In the reciprocal assessment, where T3D^TD^-derived genes were mono-reassorted into a PL^RG^ background, M1, S4 and L3 also individually reduced plaque size relative to parental PL^RG^ (Figure 2E). Given that these three genes were incapable of fully restoring the large plaque size of PL-derived T3D strain, we tested the genes in combinations of 2 or 3. The M1 gene had the greatest impact on plaque size, any combination of M1, S4, and L3 provided additive enhancement of plaque size relative to independent genes. Ultimately, the combination of all three genes could restore the parental phenotype. Our genetic evaluation therefore implicated S4, M1 and L3 genes as having a pronounced impact on replication of T3D^PL^ versus T3D^TD^.

Reovirus particles are composed of two concentric proteinaceous capsid layers (Figure 3A). The inner capsid (core) consists of proteins σ2, λ1, and λ2, while the outer capsid, which is removed during entry, consists of μ1C, σ3, and σ1. The S4, M1, and L3 genome segments encode viral proteins σ3, μ2, and λ1 proteins, respectively. More comprehensive descriptions of these genes will follow during discovery of their specific contribution to oncolytic differences among T3D lab strains. As a general description, σ3 fulfills several known functions; it is the outermost capsid protein, contributes to stability of reovirus particles (Wetzel et al., 1997; Wilson et al., 2002), and must be removed by proteolysis during reovirus entry to reveal the transcriptionally-active core particles that establish infection (Acs et al., 1971; Chang and Zweerink, 1971; Skehel and Joklik, 1969). In addition, σ3 binds and sequesters dsRNA from cellular sensors of pathogen associated molecular patters (PAMPS) (Huismans and Joklik, 1976; Imani and Jacobs, 1988; Lloyd and Shatkin, 1992). Reovirus μ2 serves at least two functions known functions; within the virion core, it provides NTPase activity and serves as a co-factor for the viral RNA-dependent RNA polymerase λ3 (Kim et al., 2004; Noble and Nibert, 1997b). In addition, μ2 bridges viral proteins to tubulin to promote formation of localized sites of virus replication called factories. Specifically μ2 links tubulin to the viral non-structural protein μNS, which then associates with viral structural proteins and the ssRNA-binding σNS (Broering et al., 2004; Broering et al., 2002; Parker et al., 2002). The primary role of λ1 is to form the inner core of reovirus particles, though additional functions have been suggested (Bisaillon et al., 1997; Lemay and Danis, 1994; Noble and Nibert, 1997a). The multi-functionality of viral proteins makes it challenging to infer precise mechanisms for amino acid variations, and therefore an in-depth comparison of T3D^PL^, T3D^KC^, and T3D^TD^ in terms of virus particle composition and efficiency of completing the various steps of the viral replication cycle was necessary.

**Figure 3.**
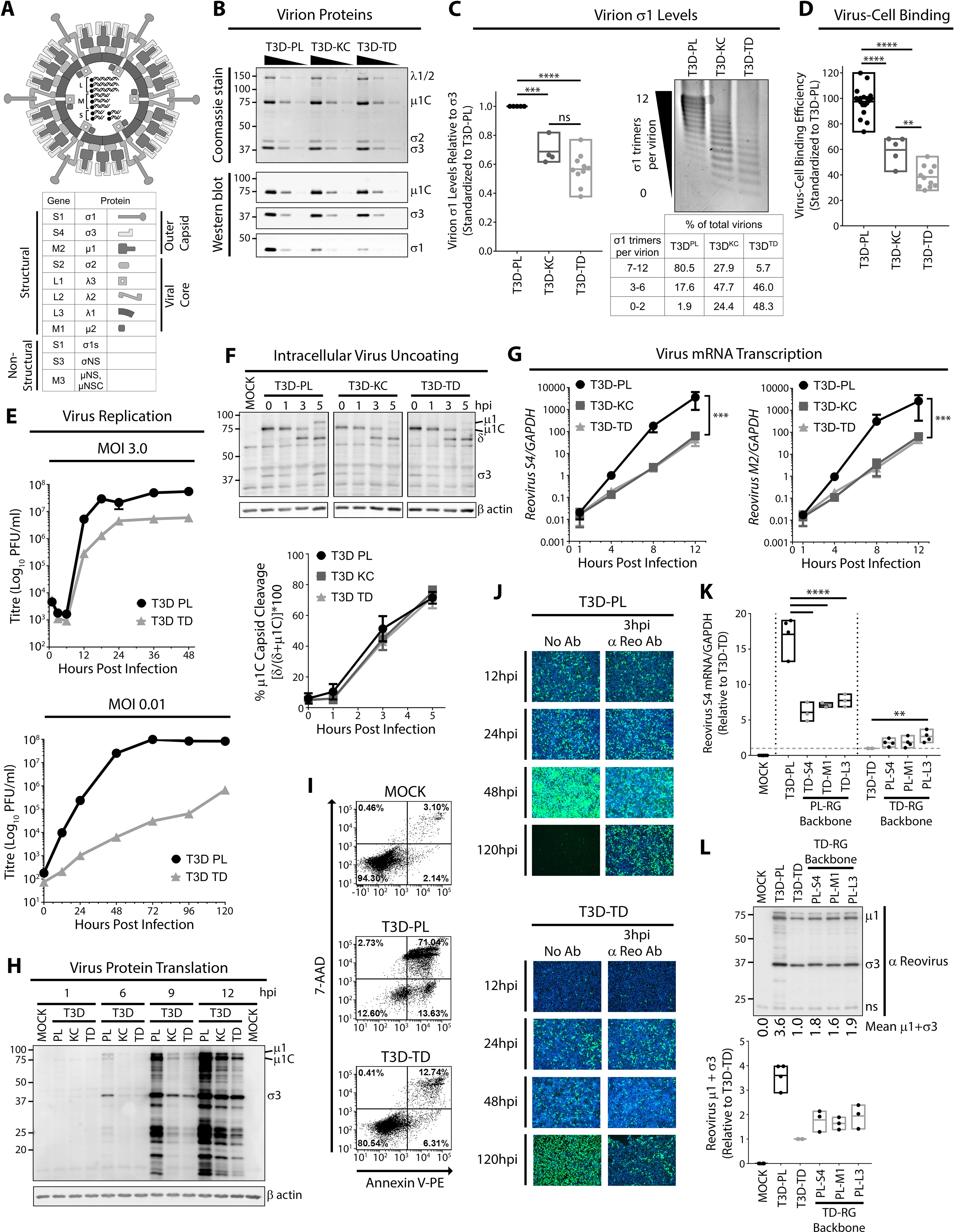
T3D^PL^ replicates more rapidly and to higher burst size in a single round of infection. **(A)** Diagrammatic representation of reovirus virion. Outer capsid proteins include σ1, σ3 and μ1. Inner capsid (core) proteins include λ2, σ2, λ1. Within the viral core are λ3, μ2 and 10 segments of dsRNA genome. **(B)** Levels of reovirus proteins were assessed using CsCl-purified virus preparations separated by SDS PAGE. Total protein was visualized by Coomassie dye staining whereas specific reovirus proteins were identified using Western blot analysis with reovirus protein specific antibodies, as indicated. **(C)** (Left) Densitometric band quantification from of σ1 relative to σ3. n ≥ 4. Statistical significance determined using one-way ANOVA with Tukey’s multiple comparisons test, *** p < 0.001, **** p < 0.0001, ns > 0.05. (Right) Agarose gel separation CsCl-purified virus preparations and based on σ1 trimers per virion and stained with Coomassie dye. Table below indicates percentage of virions with specific σ1 timers/virion calculated using densitometric band quantification. **(D)** T3D^PL^, T3D^KC^ and T3D^TD^ dilutions were bound at 4°C to non-enzymatically detached L929 cells in suspension, and following extensive washing to remove unbound reovirus, cell-bound reovirus was stained using polyclonal reovirus antibodies and mean fluorescence intensity (MFI) quantified using flow cytometry. Following T3D^PL^ standard curve generation linear regression analysis, relative cell binding of T3D^KC^ and T3D^TD^ was determined. n ≥ 5. Statistical significance determined using one-way ANOVA with Tukey's multiple comparisons test, ** p < 0.01, **** p < 0.0001. **(E)** L929 cells were infected with T3D^PL^ or T3D^TD^ at MOI 3 (left) or MOI 0.01 (right) and incubated at 37°C. At each timepoint, total cell lysates were harvested, and virus titres performed in duplicate on L929 cells. **(F)** T3D^PL^, T3D^KC^ and T3D^TD^ (MOI 1-3) were bound to L929 cell monolayers at 4°C and following washing to remove unbound virions, cells were incubated at 37°C and cell lysates were collected at various timepoints post infection. (Top) Total proteins were separated using SDS PAGE and Western blot analysis with specific antibodies were used to identify reovirus proteins and β actin (loading control). (Bottom) Densitometric band quantification of μ1C and δ and % virus uncoating was calculated using the formula [δ/(μ1C+ δ)]*100. n = 4 ± SD (MOI 1 n=2, MOI 3 n=2), linear regression analysis determined the slopes are not significantly different. **(G)** and **(H)** Similar experimental outline to (F) (MOI 3) except cell lysates were collected for RNA extraction (G) or total protein analysis (H) at indicated timepoints. (G) Total RNA was extracted, converted to cDNA and reovirus S4 and M2 RNA expression relative to housekeeping gene GAPDH was quantified using qRT-PCR. n = 3 ± SD, *** p < 0.0001, linear regression analysis determined that i) slope of T3D^PL^ differs from T3D^KC^ or T3D^TD^ and ii) slope of T3D^KC^ does not differs from T3D^TD^. (H) Total proteins were separated using SDS PAGE and Western blot analysis with specific antibodies were used to identify reovirus proteins and β actin (loading control). **(I)** Following infection similar to (F), at 24 hpi, adherent and detached cells were collected and subjected to staining with 7-AAD and Annexin V, followed by flow cytometric analysis. **(J)** L929 cells were infected with T3D^PL^ or T3D^TD^ (MOI 1) and incubated at 37°C. Polyclonal reovirus antibody was added directly to the well media at 3hpi (3hpi αReo Ab). Reovirus infected cells were visualized by fluorescence microscopy using polyclonal anti-reovirus antibodies (green) and HOECHST staining of cell nuclei (blue). **(K)** Standardized for equal infection (MOI 3), L929 cells were infected with parental T3D^PL^ or T3D^TD^, and S4, M1 and L3 gene monoreassortant PL-RG or TD-RG viruses. At 12hpi, total RNA was extracted, converted to cDNA and reovirus S4 RNA expression relative to housekeeping gene GAPDH was quantified using qRT-PCR. n ≥ 3. All values were normalized to T3D^TD^. Statistical significance determined using one-way ANOVA with Dunnett's multiple comparisons test, * p < 0.05, *** p < 0.001, **** p < 0.0001, ns > 0.05. **(L)** Similar experimental outline to (K) except only the parental T3D^PL^ or T3D^TD^, and S4, M1 and L3 gene monoreassortant TD-RG viruses were assessed at 12hpi. (Left) Total cell lysates were collected at 12hpi for Western blot analysis using specific antibodies to identify reovirus proteins. (Right) Densitometric band quantification of μ1 and σ3, relative to non-specific background band. n ≥ 3.

### Enhanced attachment of T3D^PL^ virions to cells due to increased numbers of σ1 trimers

Previous studies demonstrated that modifications to outer capsid proteins such as σ1, σ3 and μ1C can impact reovirus infectivity (Mohamed et al., 2015; Shmulevitz et al., 2012). For example, levels of the σ1 cell attachment proteins were previously found to affect cell binding (Larson et al., 1994; Mohamed et al., 2015), while mutations in σ3 and μ1 could affect the efficiency of virus uncoating (Danthi et al., 2008; Doyle et al., 2012). To determine if T3D laboratory strains exhibit distinct protein compositions, CsCl gradient-purified T3D^PL^, T3D^KC^ and T3D^TD^ virus particles were subjected to denaturing gel electrophoresis and proteins visualized by Imperial^TM^ total protein staining (Figure 3B top) and Western blot analysis using reovirus protein specific antibodies (Figure 3B bottom). The ratio of capsid proteins λ1/2, μ1C, σ2 and σ3 were similar between T3D^PL^, T3D^KC^ and T3D^TD^. However, for equivalent capsid protein (σ3), T3D^KC^ and T3D^TD^ had lower average levels of σ1 compared to T3D^PL^ (Figure 3B bottom, 3C left).

The σ1 protein is a trimer that extends from channels generated by pentameric λ2 proteins at vertices of reovirus particles, and functions as the cell-attachment protein. Not all reovirus virion vertices need to be occupied by σ1; in fact, reovirus particles can have 0-to-12 σ1 trimers depending on the sequence of σ1 or λ2 (Mohamed et al., 2015; Shmulevitz et al., 2012). We and others previously observed that 3 or more σ1 trimers-per-virion are necessary and sufficient for virion binding to cells (Larson et al., 1994; Mohamed et al., 2015). Given the importance of having ≥3 σ1 trimers for attachment, we characterized the number of σ1 trimers-per-virion for each T3D reovirus laboratory strain. Since Western blot analysis provides only an average level of virion σ1, agarose gel separation of full virions was applied as a method to distinguish virions based on σ1-per-virion levels (Figure 3C right). More than 80% of T3D^PL^ particles possessed a range of 7-12 σ1 trimers-per-virion. Contrastingly, only 28% and 6% of T3D^KC^ and T3D^TD^ particles, respectively, had 7-12 σ1 trimers-per-virion. Close to 50% of T3D^KC^ and T3D^TD^ virions had a range of 3-6 σ1 trimers/virion. T3D^KC^ and T3D^TD^ had 24% and 48% of virions with 0-2 σ1-per-virion, respectively. Most-importantly, 24% and 48% of T3D^KC^ and T3D^TD^ particles (respectively), but only 2% of T3D^PL^ particles, had fewer than 3 σ1 trimers-per-virion.

Since a greater proportion of T3D^PL^ particles had more than 3 σ1 trimers-per-virion, we predicted that a greater proportion of T3D^PL^ particles would successfully attach to cells. To compare cell attachment efficiency among the three laboratory strains, L929 cells were exposed to equivalent particle numbers of each laboratory strain at 4°C for 1 hour to permit virus binding without entry, then washed extensively. Flow cytometric analysis was used to measure bound virions per cell, reflected by the mean-fluorescence intensity (Figure 3D). Flow cytometry provides a highly linear assessment of reovirus binding over a wide range (81- fold) of virus dilutions (Supplementary Figure 1). T3D^KC^ and T3D^TD^ bound cells at 60% and 40% efficiency of T3D^PL^, respectively, which correlated closely to the percentage of particles with fewer than 3 σ1 trimers-per-virion. Together our analysis indicates that oncolytic potency of the most-oncolytic laboratory strain T3D^PL^ correlates with enhanced attachment to tumor cells and higher proportion of virus particles with ≥3 σ1 trimers-per-virion. Accordingly, assessment of σ1 levels-per-virion could be used as one of the measures of good binding and oncolysis.

The σ1 proteins of T3D^TD^ and T3D^KC^ differ from T3D^PL^ at 2 positions (Figure 1I). We consider it likely that the alanine-to-valine change at position 22 is responsible for decreased levels of σ1 on T3D^TD^ and T3D^KC^ virions, as we and others previously found that mutations in the σ1 anchor domain (amino acids 1-27) affect incorporation of σ1 trimers into the λ2 vertex (Bokiej et al., 2012; Mohamed et al., 2015; Nygaard et al., 2013). The sequence difference at residue 408 is located in the σ1 head domain (amino acids 311-455), which interacts with cell surface JAM-A (Barton et al., 2001; Campbell et al., 2005). This difference could contribute to the differential virion cell binding levels; though we think this unlikely because the change is not located at the JAM-A binding interface and because binding deficiency of T3D^KC^ and T3D^TD^ very closely correlated with having virions with less than 3 σ1 trimers (Figure 3C versus 3D).

Importantly, the S1 genome segment that encodes σ1 did not strongly segregate with the large plaque phenotype in our reassortant studies (Figure 2B, 2D, 2E), likely because differences in cell binding are only ~2-fold, and therefore less impactful than contributions by M1, S4, and L3. The remainder of our studies therefore focused on understanding the mechanisms by which the M1, S4, and L3 genome segments contribute dominantly towards oncolytic activity of T3D^PL^.

### T3D^PL^ replicates more rapidly and to higher burst size in a single round of infection

The larger plaques produced by T3D^PL^ could stem from more rapid rates of virus replication, increased burst size, enhanced cell lysis and virus release, or increased cell-to-cell spread. To determine if T3D^PL^ replicates more rapidly and/or to higher burst size than T3D^TD^ in a single round of infection, single-step virus growth analyses were performed at a multiplicity of infection (MOI) of 3 (Figure 3E top). L929 cells were exposed to T3D^PL^ and T3D^TD^ at 4°C for 1 hour to synchronize infection, then incubated at 37°C for various time-points. Reovirus titres at 1 hour post infection (hpi) were similar between T3D^PL^ and T3D^TD^, indicating equal input virion titers. Following entry, reovirions uncoat to cores and become non-infectious, leading to a drop in titers at 3 and 6hpi for both T3D^PL^ and T3D^TD^. Between 8-12hpi, ~1000-fold new infectious progeny were produced per input of T3D^PL^. In contrast, titers increased by only ~50-fold for T3D^TD^. T3D^PL^ reached maximum saturation titers at 18hpi, while T3D^TD^ titers saturated at 24hpi. Furthermore, saturation titers for T3D^PL^ were ~9-fold higher than T3D^TD^. The one-step growth curves therefore indicated that new progeny production for T3D^PL^ occurs faster, and produces higher virus burst size, relative to T3D^TD^. Importantly, these findings suggest that oncolytic potential correlates with proficiency of virus replication in a single round of replication. When multi-step virus growth curves were performed at an MOI of 0.01, the cumulative advantage of improved replication over several rounds of infection was striking; T3D^PL^ accumulated ~4000-fold higher virus titers relative to T3D^TD^ by 48hpi (Figure 3E bottom).

Increased virus replication could result from higher efficiency of virus entry, or post-entry steps of virus replication and assembly. Following attachment to cells, reovirus undergoes endocytosis and trafficking to endo-lysosomes where cathepsins L and B mediate virus uncoating (Ebert et al., 2002; Johnson et al., 2009). Here, the outermost σ3 protein is degraded and μ1C is cleaved to the membrane-destabilizing products δ, μ1N and φ (Odegard et al., 2004; Zhang et al., 2006). The resulting intermediate subviral particles (ISVPs) are capable of penetrating membranes and delivering reovirus core particles to the cytoplasm. Given that efficient reovirus uncoating is a crucial step for successful infection, we compared uncoating among T3D laboratory strains. Note that in all subsequent studies, to overcome differences in cell binding, virus doses were adjusted to give equivalent cell-bound virus particles. To monitor uncoating, cleavage of σ3 and μ1C was followed by Western blot analysis at various times post-infection (Figure 3F). Complete cleavage of σ3 was observed by 3hpi and coincided with initiation of μ1C to δ cleavage. The rate of μ1C cleavage to δ was quantified as a percentage of δ to δ + μ1C, and standardized to β actin housekeeping protein. All T3D strains displayed similar uncoating rates and almost complete uncoating was attained by 5hpi. Unlike T3D^KC^ and T3D^TD^, *de-novo* synthesized μ1 (precursor to μ1C) and σ3 was observed at 5hpi in T3D^PL^, suggesting that a post-uncoating step was enhanced during T3D^PL^ infection.

Following uncoating of the outer capsid, reovirus cores that enter the cytoplasm become transcriptionally active. Positive-sense mRNAs are transcribed within reovirus cores and released into the cytoplasm for translation by host protein synthesis machinery (Acs et al., 1971; Chang and Zweerink, 1971; Skehel and Joklik, 1969). To assess core transcriptase activity and overall accumulation of viral RNA, we used quantitative RT-PCR for viral RNAs over the course of infection (Figure 3G). S4 and M2 viral RNAs were used as representative viral genes, while GAPDH was used for standardization between samples. Over 12hpi, viral transcripts accumulated at a significantly faster rate for T3D^PL^ than T3D^KC^/T3D^TD^. Specifically, prior to viral RNA saturation at 8hpi, T3D^PL^ exhibited 77- and 85-fold higher levels of S4 viral RNA compared to T3D^KC^ and T3D^TD^, respectively (Figure 3G left). Differences in viral M2 RNA were even greater, with T3D^PL^ having 92- and 133-fold higher levels than T3D^KC^ and T3D^TD^, respectively (Figure 3G right). As anticipated, viral protein levels closely reflected virus transcript levels; Western blot analysis with polyclonal anti-reovirus antibody showed substantially higher accumulation of T3D^PL^ viral proteins compared to T3D^KC^ and T3D^TD^ at every timepoint from 6hpi onwards (Figure 3H). Not surprisingly given the large increase in kinetics and total level of viral RNA and proteins for T3D^PL^, cell death measured by flow cytometry with Annexin V (measures early apoptosis) or 7-Aminoactinomycin (measures cell death) was higher for T3D^PL^ relative to T3D^TD^; for example ~85% versus 19% at MOI of 3 for these strains respectively (Figure 3I). To monitor cell-cell spread, we use immunofluorescence staining of reovirus-antigen-positive L929 cells (Figure 3J). In order to decipher the extent of cell-cell spread versus initially-infected cells, staining was compared in the absence or presence of reovirus neutralizing antibodies added at 3hpi to prevent cell-cell spread. Cells became positive for T3D^PL^ staining more rapidly than T3D^TD^ (i.e. compare viruses at 12hpi) and equalized at ~24hpi, reflecting the enhanced kinetics of RNA and protein synthesis during T3D^PL^ infection. By 48hpi, T3D^PL^ very effectively spread to neighbouring cells (i.e. compare 48hpi No Ab to anti-reovirus Ab), while limited cell-cell spread was observed for T3D^TD^. By 120hpi, very few cells remained alive following T3D^PL^ infection, indicated by a lack of Hoechst nuclear staining. In comparison, T3D^TD^ successfully underwent cell-cell spread by 120hpi but cells remained relatively intact (Figure 3J). Altogether, the analysis indicates that kinetics of reovirus macromolecule synthesis, cell death, and cell-cell spread are drastically faster for T3D^PL^ relative to T3D^TD^.

M1, S4, and L3 reovirus genes were associated with increased plaque size of T3D^PL^ (Figure 2), so we determined if these three genes also affect the levels of viral RNA and protein in the first round of infection. RNA and protein analysis were chosen over later steps in virus replication, in order to capture the earliest difference between T3D laboratory strains. Quantitative RT-PCR (Figure 3K) and Western blot analysis (Figure 3L) was repeated for single mono-reassortants at 12hpi. M1, S4, and L3 genes from T3D^PL^ increased reovirus RNAs in an otherwise TD^RG^ background, relative to T3D^TD^, while M1, S4, and L3 genes from T3D^TD^ decreased reovirus RNAs in an otherwise PL background relative to T3D^PL^. In a T3D^TD^ background, it was evident that M1, S4, and L3 genes from T3D^PL^ could increase virus protein (μ1C) levels. Importantly, when introduced independently, each gene only conferred intermediate levels of RNA and protein relative to the parental strains, indicating that each gene only confers a portion of the replication advantage that T3D^PL^ exhibits over T3D^TD^. The data therefore indicated that S4, M1 and L3 each contribute to the higher accumulation of virus macromolecules for the most-oncolytic T3D^PL^ strain relative to the less-oncolytic T3D^TD^ strain.

### M1-encoded μ2 confers accelerated core transcription

The observation that virus transcripts accumulate at a faster rate for T3D^PL^ does not directly indicate whether T3D^PL^ cores transcribe more efficiently, since reovirus RNAs are packaged into new progeny cores that amplify RNA and protein synthesis. In other words, increased transcription or translation would produce more cores and therefore more RNA and protein in subsequent rounds of amplification; therefore making it difficult to distinguish which process is directly enhanced. To directly assess transcription activities of T3D laboratory strains, we therefore performed *in-vitro* transcription assays (Figure 4A). Reovirus particles become transcriptionally active once the outer capsid proteins are uncoated to generate virus cores with only the inner-capsids intact (Acs et al., 1971; Chang and Zweerink, 1971). As is a routine approach to generate cores *in-vitro*, T3D^PL^, T3D^KC^ and T3D^TD^ particles were treated with chymotrypsin (CHT) and purified by high speed centrifugation. The purity of viral cores was confirmed by Coomassie staining and Western blot analysis (Figure 4B); cores had the characteristic loss of outer capsid proteins (σ1, σ3 and μ1) but fully retained core inner-capsid proteins (σ2, λ1, λ2). Cores were then subjected to transcription reactions for various time intervals, and levels of S4 and M2 viral RNAs measured by qRT-PCR (Figure 4C). Each sample was spiked with *in-vitro* synthesized mouse GAPDH RNA prior to RNA extraction to control for sample-to-sample processing variations. Negative control reactions lacking rATP produced no increases in RNA levels, confirming that RNA measured in the presence of all four rNTPs accurately reflected *de-novo* core transcription (Figure 4D). Furthermore, *in-vitro* core transcription rates were linear, unlike the exponential rates observed during intracellular infection, reflecting the absence of secondary rounds of amplification that occur during reovirus infection but not *in vitro*. Importantly, the rates of both S4 and M2 viral RNA transcription were significantly higher for T3D^PL^ compared to T3D^KC^ and T3D^TD^. For example S4 transcripts increased for T3D^PL^ at 16.8±0.9/minute while at 3.6±0.3 and 2.5±0.3/minute for T3D^KC^ and T3D^TD^ respectively. For M2, rates were 7.3±0.5, 1.1±0.2, and 0.9±0.1 transcripts/minute for T3D^PL^, T3D^KC^ and T3D^TD^ respectively. Therefore, T3D^PL^ has an inherent transcriptase advantage within the viral cores.

**Figure 4.**
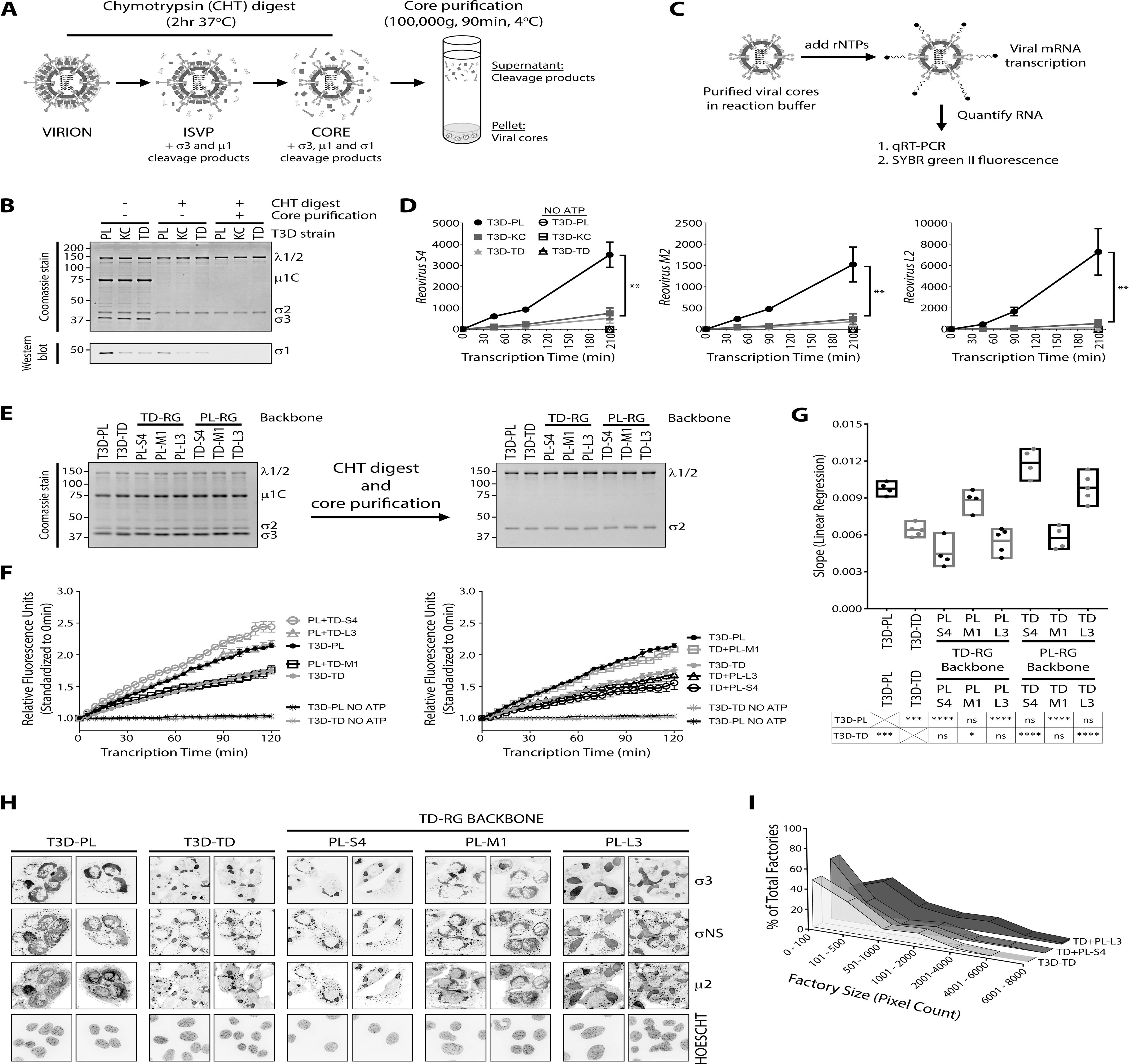
M1-encoded μ2 of T3D^PL^ confers accelerates core transcription and globular factory morphology while L3-encoded λ1 confers increased factory size. **(A)** CsCl purified preparations of T3D^PL^, T3D^KC^ and T3D^TD^ were each incubated with chymotrypsin at 37°C for 2hrs. Viral cores were purified by ultra-centrifugation at 100,000g for 90min at 4C. The pellet fraction containing viral cores was resuspended in 100mM Tris pH 8.0 and assessed for purity followed by core transcription assays. **(B)** At every stage in viral core purification, aliquots were separated by SDS-PAGE and total protein was visualized with Coomassie dye, while reovirus σ1 protein was identified using Western blot analysis with σ1-specific antibody. **(C)** Purified viral cores from A) were added to transcription buffer (with or without ATP) and incubated at 40°C. 1) Each timepoint sample was spiked with 3ng mouse GAPDH RNA. Following total RNA extraction using Trizol LS and cDNA synthesis, qRT-PCR was performed using gene-specific primers for reovirus S4, M2, L2 or GAPDH. 2) Transcription reactions were spiked with SYBR green II RNA dye and fluorescence was monitored at 5min intervals. **(D)** Following core transcription reaction as in C1), reovirus RNA (S4, M2, L2) values were standardized to respective GAPDH, and each group was normalized to the respective 0min (input) timepoint. n = 3 ± SD, ** p < 0.001, linear regression analysis determined that i) slope of T3D^PL^ differs from T3D^KC^ or T3D^TD^ and ii) slope of T3D^KC^ does not differs from T3D^TD^. **(E)** Viral cores were purified as described in (A) from parental T3D^PL^ or T3D^TD^, and S4, M1 and L3 gene monoreassortant PL-RG or TD-RG viruses. Aliquots were separated by SDS-PAGE and total protein was visualized with Coomassie dye, prior to CHT incubation (Left) and viral core purification (Right). **(F)** Core transcription reaction were set-up as in C2), and fluorescence values was standardized to 0min (input) timepoint. Parental T3D^PL^ or T3D^TD^, and S4, M1 and L3 gene monoreassortant PL-RG (Left) or TD-RG (Right) viruses. n ≥ 4 ± SEM. **(G)** Linear regression analysis performed on data from (F). n ≥ 4, Statistical significance determined using one-way ANOVA with Tukey's multiple comparisons test, with either parental T3D^PL^ or T3D^TD^ as control dataset. * p < 0.05, *** p < 0.001, **** p < 0.0001, ns > 0.05. **(H)** ID8 cells seeded on coverslips were infected (MOI 3) with parental T3D^PL^ or T3D^TD^, and S4, M1 and L3 gene monoreassortants in a TD-RG backbone, for 20hpi. Cells were fixed and stained with reovirus protein specific antibodies (σNS, μ2, σ3). Nuclei were stained with HOESCHT 33342. Cells were imaged using confocal microscopy. Projection images from two fields of view for each condition are shown. **(I)** Using σNS images from (H), reovirus factory size was quantified using ImageQuant colony counting software and % of total factories was determined for indicated factory size range. Viral factories were analyzed for 4-5 fields of view, and total viral factories assessed were T3D^TD^ 195, TD+PL-S4 193, TD+PL-L3 203.

Having identified core transcription as a key mechanism for the superior replication T3D^PL^, we then asked if any of the 3 genes that segregated with large-plaque phenotype of T3D^PL^ (Figure 3E-G) account for the inherent core transcription advantage. The M1-encoded μ2 protein seemed a likely candidate, as one of the μ2 functions is to hydrolyze NTPs (NTPase) and act as a co-factor to the RNA-dependent RNA polymerase λ3 (Kim et al., 2004; Noble and Nibert, 1997b). The inner-capsid λ1 protein encoded by L3 could indirectly impact the structure and activity of cores, while the σ3 viral protein encoded by the S4 gene is completely removed from cores and therefore was not anticipated to affect core transcription *in vitro*. Using parental and mono-reassortant viruses generated in Figure 2, we assessed the effects of M1, S4, and L3 genes on core transcription. All viruses were successfully uncoated to cores with chymotrypsin digestion (Figure 4E) and subjected to *in vitro* transcription reactions (Figure 4F). To facilitate more-rapid comparison among many samples, we developed an assay that measures RNA levels in real-time throughout the reaction, using RNA-binding SYBR II. It should be noted that the SYBR II method has notably lower detection sensitivity than qRT-PCR (i.e. compare readouts for Figure 4F versus 4D), but nevertheless consistently recapitulated the enhanced transcriptional rates of T3D^PL^ relative to T3D^TD^, and was chosen over qRT-PCR because it eliminated the cumbersome steps of spike-in controls, RNA purification and qRT-PCR for every time point. As expected, no RNA was produced in the absence of rATP (negative control). When the T3D^PL^-derived M1 was introduced into an otherwise T3D^TD^ background, it was sufficient to restore core transcription to rates similar to T3D^PL^ (Figure 4F right). Reciprocally, T3D^TD^-derived M1 decreased RNA synthesis rates when placed into an otherwise T3D^PL^ background, relative to T3D^PL^ (Figure 4F left). When slopes of transcription rates were graphed for three independent experiments (Figure 4G), it was evident that the M1 gene was necessary and sufficient to confer high transcription rates of T3D^PL^. These data clearly demonstrate that the M1-encoded μ2 protein contributes to improved transcription by T3D^PL^, contributing to the positive impact of μ2 on the oncolytic potency of T3D^PL^.

### M1-encoded μ2 and L3-encoded λ1 alter virus factory morphology and increase factory size respectively

A second characterized function of M1-encoded μ2 is to help organize newly made viral proteins at localized sites of virus replication called “factories” (Broering et al., 2004; Broering et al., 2000a; Broering et al., 2002; Parker et al., 2002). Shortly after escape of the endosome into the cytoplasm, reovirus cores establish factories by localized production of viral RNAs and proteins, and an active strategy to recruit newly synthesized viral proteins (Broering et al., 2004; Broering et al., 2000a; Broering et al., 2002). Specifically, three reovirus proteins play an essential role in factory formation: the non-structural protein μNS recruits *de-novo* synthesized core proteins (λ1, λ2, λ3, σ2, μ2) (Becker et al., 2003; Broering et al., 2005; Broering et al., 2004; Broering et al., 2000b; Broering et al., 2002; Miller et al., 2007; Miller et al., 2010; Miller et al., 2003), σNS binds RNAs, and μ2 tethers this complex to the cytoskeleton (Broering et al., 2004; Broering et al., 2000a; Broering et al., 2002; Miller et al., 2003). Host protein translational machinery is also recruited to the periphery of factories through unknown mechanisms (Desmet et al., 2014). One of the amino acid differences in μ2 between T3D^PL^ and T3D^TD^ is at position 208, which was previously demonstrated to affect virus factory morphology by comparison of serotype 3 versus 1 reoviruses (Kobayashi et al., 2009; Yin et al., 2004). Specifically, a serine at position 208 in T3D^TD^ was associated with a globular factory morphology, while a proline at the same position caused more tubulin-associated fibrous morphology (Kobayashi et al., 2009; Yin et al., 2004). Studies have yet to demonstrate a relationship between factory characteristics and effects on virus titers. Most-recently however, factory morphology was found to affect the encapsidation efficiency of genomic RNAs, as globular factories were associated with an increased fraction of genome-devoid (empty) virions (Shah et al., 2017).

To determine if T3D laboratory strains differed in factory morphology, immunofluorescence staining was conducted on T3D^PL^ and T3D^TD^ infected L929 cells. Reovirus factories were detectable by staining for factory-associated proteins σNS and μ2, and also the outercapsid protein σ3 (Figure 4H). Not surprisingly, the T3D^PL^ which has a proline at position 208 of μ2 similar to T1L, showed fibrous morphology. As anticipated, substitution of the T3D^PL^-encoded M1 into an otherwise T3D^TD^ genomic background, recapitulated the fibrous factory morphology of T3D^PL^. Our analysis therefore shows that the molecular basis for enhanced *in vitro* replication and oncolysis of T3D^PL^ includes an increase in transcription kinetics conferred by μ2 (Figure 4A-G), but perhaps also relates to μ2-dependent establishment of tubulin-associated fibrous viral factories (Figure 4H).

The most striking and unexpected result of immunofluorescence staining of reovirus factories was that L3-encoded λ1 was clearly associated with establishing factory size (Figure 4H, quantified in Figure 4I). As mentioned, the λ1 protein is an essential component of the reovirus core, which is composed of λ1-λ1 homodimers and λ1-σ2 heterodimers. Amino acid differences between λ1 T3D^PL^ and T3D^TD^ laboratory strains are at λ1-λ1 and λ1-σ2 interfaces, as will be further depicted and described in the discussion. While the T3D^PL^-derived L3/λ1 in an otherwise T3D^TD^ genomic background did not affect the morphology of factories (i.e. they remained globular), it starkly increased the size of the globular factories (Figure 4I). Keeping in mind that M1 and S4 monoreassortants had similar levels of viral RNA and protein as L3, yet did not increase factory size, the findings suggest a novel role for λ1 in accumulation of viral macromolecules efficiently at virus factories. This is the first time, to our knowledge, that differences in factory size (rather than morphology) are described between reoviruses and correlated with improved virus amplification.

### T3D^TD^ activates interferon signaling more than T3D^PL^, but IFN signaling does not impact the first round of reovirus infection

In addition to virus factors that impact virus replication, host interferon signaling can inhibit virus infection. Upon entry into host cells, RNA viruses activate pathogen associated molecular pattern (PAMP)-recognition receptors such as cytosolic retinoic acid-inducible gene I (RIG-I), melanoma differentiation-associated factor 5 (MDA5), laboratory of genetics and physiology 2 (LGP2)), protein kinase R (PKR) and endosomal toll-like receptor 3 (TLR3). Although reovirus dsRNA genomes are encased in core particles, it is proposed that reovirus PAMPs are generated from occasional unstable cores and/or viral mRNA secondary structures (Henderson and Joklik, 1978). Activation of PAMP receptors, through a cascade of adaptor proteins and signaling events, results in phosphorylation (activation) and nuclear translocation of IRF3/7 and NFκB transcription factors, which subsequently induce expression of interferons (IFNs), antiviral interferon stimulated genes (ISGs) and inflammatory cytokines (Figure 5A). Numerous studies have demonstrated the importance of the RIG-I/MDA5 signaling axis on reovirus mediated IFN production and subsequent paracrine suppression of reovirus spread to neighbouring cells (Goubau et al., 2014; Loo et al., 2008; Shmulevitz et al., 2010b). However, whether autocrine RIG-I/MDA5 and IFN signaling can reduce the initial round of reovirus infection is less clearly understood. Therefore an obvious question was whether T3D^TD^ induces more robust IFN signalling, and if so, whether this contributes to the reduced replication of T3D^TD^ relative to T3D^PL^.

**Figure 5.**
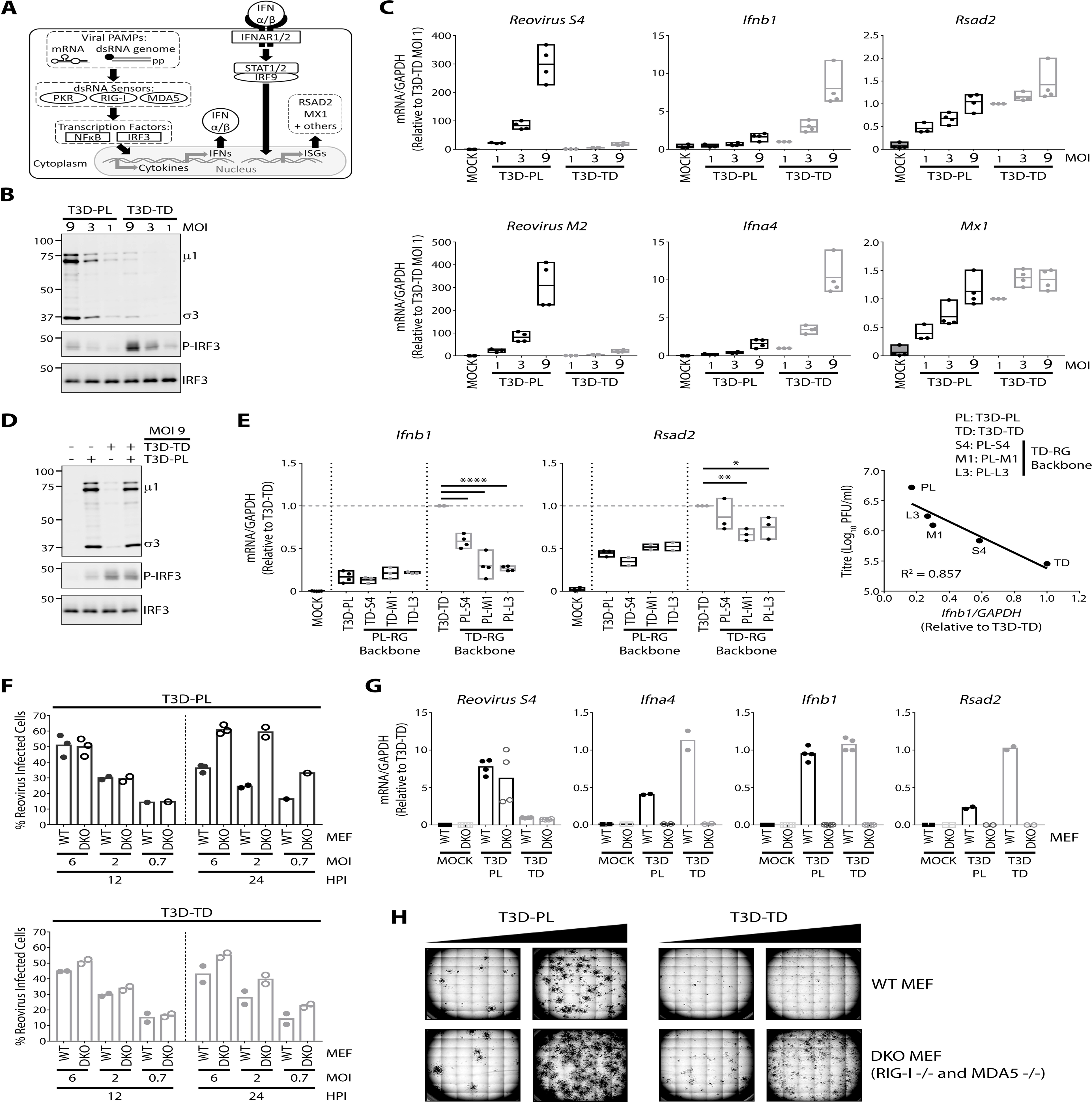
T3D^TD^ activates interferon signaling more than T3D^PL^, but IFN signaling does not impact the first round of reovirus infection. **(A)** Overview of antiviral signaling induced during reovirus infection. **(B-D)** L929 cells were infected with T3D^PL^ and/or T3D^TD^ at indicated MOI and incubated at 37°C for 12 hours. (B) and (D) Total proteins were separated using SDS PAGE and Western blot analysis with indicated antibodies. C) Total RNA was extracted, converted to cDNA and gene expression relative to housekeeping gene GAPDH was quantified using qRT-PCR. All values were normalized to T3D^TD^ MOI 1. n = 4. **(E)** Standardized for equal infection (MOI 3), L929 cells were infected with parental T3D^PL^ or T3D^TD^, and S4, M1 and L3 gene monoreassortant PL-RG or TD-RG viruses. (Left) At 12hpi, total RNA was extracted, converted to cDNA and gene expression relative to housekeeping gene GAPDH was quantified using qRT-PCR. All values were normalized to T3D^TD^. n ≥ 3. Statistical significance determined using one-way ANOVA with Dunnett's multiple comparisons test, * p < 0.05, *** p < 0.001, **** p < 0.0001, ns > 0.05. (Right) At 12hpi, total viral titres at 12hpi were plotted against *Ifnb1* gene expression, followed by linear regression analysis. **(F-H)**WT or RIG-I/MDA5 -/- double knockout (DKO) MEFs were infected with T3D^PL^ or T3D^TD^. (F) At indicated MOIs and timepoints, percent reovirus infected cells were identified using reovirus specific primary antibody and Alexa Fluor 488 conjugated secondary antibody and quantified using flow cytometry. n = 1-3, (G) For MOI 6 at 12hpi, total RNA was extracted, converted to cDNA and gene expression relative to housekeeping gene GAPDH was quantified using qRT-PCR. All values were normalized to WT MEF T3D^TD^. n ≥ 1-2, with experimental duplicates, (H) At 3days post infection, reovirus infected cell foci were stained with colorimetric immunocytochemistry using primary polyclonal reovirus antibody, alkaline phosphatase secondary antibody and BCIP/NBT substrate.

First we determined if there were differences in IFN signalling between T3D laboratory strains. L929 cells were infected with T3D^TD^ or T3D^PL^ at a range of doses (MOI 1, 3, and 9), or mock infected. IFN signalling was then assessed at 12hpi, an intermediate timepoint of virus replication when the potentially confounding effects of cell-cell spread of virus are minimal. IRF3 phosphorylation (activation) was strongly induced by T3D^TD^ but not T3D^PL^, despite a reciprocal trend for reovirus protein expression (Figure 5B). Moreover, while both T3D laboratory strains caused a dose-dependent increase in transcripts of IFN (*Ifnα4*, *Ifnβ*) and IFN-inducible genes (*Mx1*, *Rsad2*), T3D^TD^ induced higher expression relative to T3D^PL^, as assessed by qRT-PCR (Figure 5C). In other words, despite producing lower levels of viral proteins and transcripts, T3D^TD^ induced elevated levels of antiviral signaling compared to T3D^PL^.

We considered two alternative explanations to account for increased IFN signalling by T3D^TD^: i) T3D^TD^ might be a more potent inducer of antiviral signaling, or ii) T3D^PL^ is a more potent inhibitor of antiviral signaling. To distinguish between these possibilities, L929 cells were co-infected with T3D^PL^ and T3D^TD^ at a high MOI of 9 (each) to ensure that most cells were infected with both viruses, and cell lysates were subjected to Western blot analysis for IRF3 phosphorylation. Phospho-IRF3 levels were similar between T3D^TD^ and T3D^PL^/T3D^TD^ co-infection (Figure 5D), suggesting that T3D^TD^-dependant activation of IRF3 could not be overcome by the presence of T3D^PL^. Therefore, T3D^TD^ is most likely a more potent activator of antiviral signaling than T3D^PL^. In other words, paradoxically, the less-prolific replicating variant is more dominant for IFN expression. Furthermore, the levels of reovirus proteins were either unchanged (Figure 5D, σ3) or only marginally reduced (Figure 5D, μ1/μ1C) in T3D^PL^/T3D^TD^ co-infection compared to T3D^PL^ despite high phospho-IRF3 levels, suggesting that IRF3 activation and downstream signaling may not play a major role in restricting the first round of reovirus replication.

Next, we determined if any of the 3 genes that segregated with the large plaque phenotype of T3D^PL^ (i.e. S4, M1, and L3) contributed to the differential activation of IFN signalling between the two virus strains, by analyzing mono-reassortants. As previously noted (Figure 3I and 3J), the mono-reassortants of S4, M1, and L3 produced intermediate viral RNA and protein levels compared to the T3D^PL^ and T3D^TD^ parental strains, reflecting their intermediate levels of replication. As for IFNs and IFN-induced genes, mono-reassortants also gave intermediate IFN signalling and no mono-reassortant fully reversed the phenotype of IFN signalling (Figure 5E). For example, IFNβ was induced more by T3D^TD^ than T3D^PL^ parental strain, but individually adding S4, M1, or L3 from T3D^TD^ into an otherwise T3D^PL^ genomic background, did not increase IFNβ levels to those achieved by T3D^TD^. One possible interpretation of this data is that S4, M1, and L3 each independently contribute ‘somewhat’ to IFN signalling, such that mono-reassortants are insufficient to confer the full parental phenotype. Previous studies have indeed implicated these genes (and other genes such as S1) in affecting IFN signalling (Beattie et al., 1995; Lanoie and Lemay, 2018; Zurney et al., 2009). Of these IFN modulating viral proteins, the S4-encoded σ3 has been clearly demonstrated to sequester dsRNA, inhibit activation of PKR and rescue other viruses depleted of inhibitors of antiviral response (Beattie et al., 1995; Denzler and Jacobs, 1994; Yue and Shatkin, 1997). But it should be noted that most studies on reovirus genes that impact IFN signalling do not consider whether effects are direct (e.g. the gene or protein directly modulate IFN mediators), or whether instead the viral genes impact virus replication and thereby indirectly impact IFN induction. To consider the differences in virus replication kinetics between parental and mono-reassortant viruses, we determined the relationship between IFN signaling (IFNβ mRNA) and measures of virus replication (progeny titers) (Figure 5E right). A strong negative correlation (R^2^=0.86) between virus replication proficiency and IFN signalling was found, suggesting that S4, M1, or L3 could contribute to differences in IFN signalling between T3D^TD^ and T3D^PL^ indirectly, by affecting the extent of virus replication. Specifically, we propose that incoming cores must establish viral RNA and protein expression, and factory formation around the core, with sufficient speed to prevent detection of foreign virus patterns by the host. Indeed, recent studies showed the ability of reovirus factories to selectively sort IFN signalling components (Stanifer et al., 2017). Our data is therefore consistent with a model where the extent of IFN signalling is inversely related to the efficiency of virus replication; this paradox will be further explored in the discussion.

Given that T3D^TD^ induced more IFN signalling than T3D^PL^, the pivotal question became whether IFN signalling contributed to reduced replication of T3D^TD^ relative to T3D^PL^. While it is well established that IFN signalling can prevent dissemination of reovirus to neighboring cells through paracrine signalling (Shmulevitz et al., 2010b), it is unknown whether IFN signalling can affect the initial infection of reovirus in an autocrine manner. To address this question, we made use of double knock-out (DKO) mouse embryo fibroblasts (MEFs) lacking both RIG-I and MDA5 (Errett et al., 2013; Loo et al., 2008). We reasoned that if IFN signalling can impact the first round of virus infection, then the DKO cells should demonstrate increased infection by reovirus at an intermediate timepoint of 12hpi where cell-cell spread is minimal. Wild-type (WT) and DKO MEFs were exposed to T3D^PL^ or T3D^TD^ at MOIs of 0.7, 2, and 6 (based on WT MEF titers) and flow cytometric analysis was conducted to measure the number of cells positive for reovirus antigen expression (Figure 5F). At 12hpi, WT and DKO cells showed equivalent infection at matched MOIs, indicating that IFN signalling likely does not affect the first round of infection. As expected, at 24hpi when reovirus already spreads to new cells, the DKO cells showed enhanced infection relative to WT, supporting the paracrine contribution of IFNs to reducing virus dissemination. To confirm the absence of IFN signalling in DKO cells, qRT-PCR was conducted at 12hpi for IFNs (*Ifnb1*, *Ifnα4*) and IFN-induced gene *Rsad2*; all demonstrated a strong inhibition (> 97%) of IFN signaling relative to WT cells following infection by either T3D^PL^ or T3D^TD^ (Figure 5G). Analysis of reovirus transcript levels by qRT-PCR confirmed that WT and DKO cells supported equal levels of virus replication during the initial round of infection.

While having minimal-to-no effect on the first round of T3D reovirus infection, IFN signalling did have the predicted activity on restricting cell-cell spread of reovirus. Specifically, when plaque assays were used to assess the overall replication and spread of T3D^PL^ and T3D^TD^, both viruses produced the same number of reovirus-infected cell foci on DKO MEFs versus wildtype MEFs (Figure 5H), suggesting that the initial round of infection was independent of IFN signaling. Plaques for both T3D^PL^ and T3D^TD^ were larger on DKO MEFs relative to wildtype MEFs, supporting the importance of IFN signalling during cell-cell spread. Importantly however, plaque size of T3D^TD^ remained much smaller than T3D^PL^ even on DKO MEFs; this supports the model that the oncolytic advantage of T3D^PL^ relative to T3D^TD^ is not dependent of IFN signalling, but rather dependent on differences in virus replication discovered in Figures 3 and 4. Similar results were obtained in the NIH/3T3 mouse fibroblast cell line in which RIG-I was knocked down using shRNA (Supplemental Figure 3). Altogether these results strongly suggest that RIG-I signaling does not affect the first round of reovirus replication for either T3D laboratory strain. However, the differences in IFN signalling appear to impact subsequent rounds of infection, permitting T3D^PL^ to disseminate more efficiently. Furthermore as explored in the discussion, the finding that T3D^TD^ induces more IFN signalling than T3D^PL^ is likely to also indirectly affect the landscape of anti-tumor and anti-viral immune cells and therefore is an impactful discovery for understanding the contribution of virus genetics on the immunotherapeutic aspect of virus oncolysis.

### T3D^PL^ S4-encoded σ3 stimulates expression of NFκB-dependent but IFN-independent cytokines

Having eliminated IFN as being relevant in the first round of infection, we turned to other signalling pathways in the cell. The MAPK/ERK, p38 stress-activated kinase, and NF-κB pathways have all been implicated in replication of an assortment of viruses (Bonjardim, 2017; Lim et al., 2016; Mohamed and McFadden, 2009; Schmitz et al., 2014). As for reovirus specifically, ERK, p38, and NF-κB signalling were positively associated with reovirus oncolytic activities (Norman et al., 2004; Shmulevitz et al., 2010b; Thirukkumaran et al., 2017). Western blot analysis was therefore conducted to monitor total levels versus phosphorylation status of ERK p42 and p44 subunits, p38, and NF-κB factor IκBα, at 12hpi following infection at MOI= 1 (Figure 6A). Whereas phosphorylation of p42/p44 and p38 are a direct indication of kinase activity, the phosphorylation of IκBα results its degradation from the NF-κB complex and facilitates NF-κB nuclear translocation and subesequent activity. Densitometric analysis for three independent experiments showed that T3D^PL^ induced higher levels of phospho-p38, phospho-ERK, and phospho-IκBα. Of the three signalling proteins assessed, only IκBα became differentially phosphorylated by T3D laboratory strains in a gene-dependent manner. Specifically, T3D^PL^ induced accumulation of phosphorylated IκBα while T3D^TD^ did not. Moreover, the phosphorylation of IκBα corresponded with the T3D^PL^-derived S4/σ3, since addition of this T3D^PL^ gene into an otherwise T3D^TD^ background was sufficient for NF-κB activation. The σ3 protein has two well characterized activities; it functions as an outer-capsid protein and it sequesters viral RNAs away from cellular dsRNA-detecting signalling molecules such as PKR and RIG-I (Denzler and Jacobs, 1994; Yue and Shatkin, 1997). Our data now suggested that σ3 may contribute to NF-κB activation.

**Figure 6.**
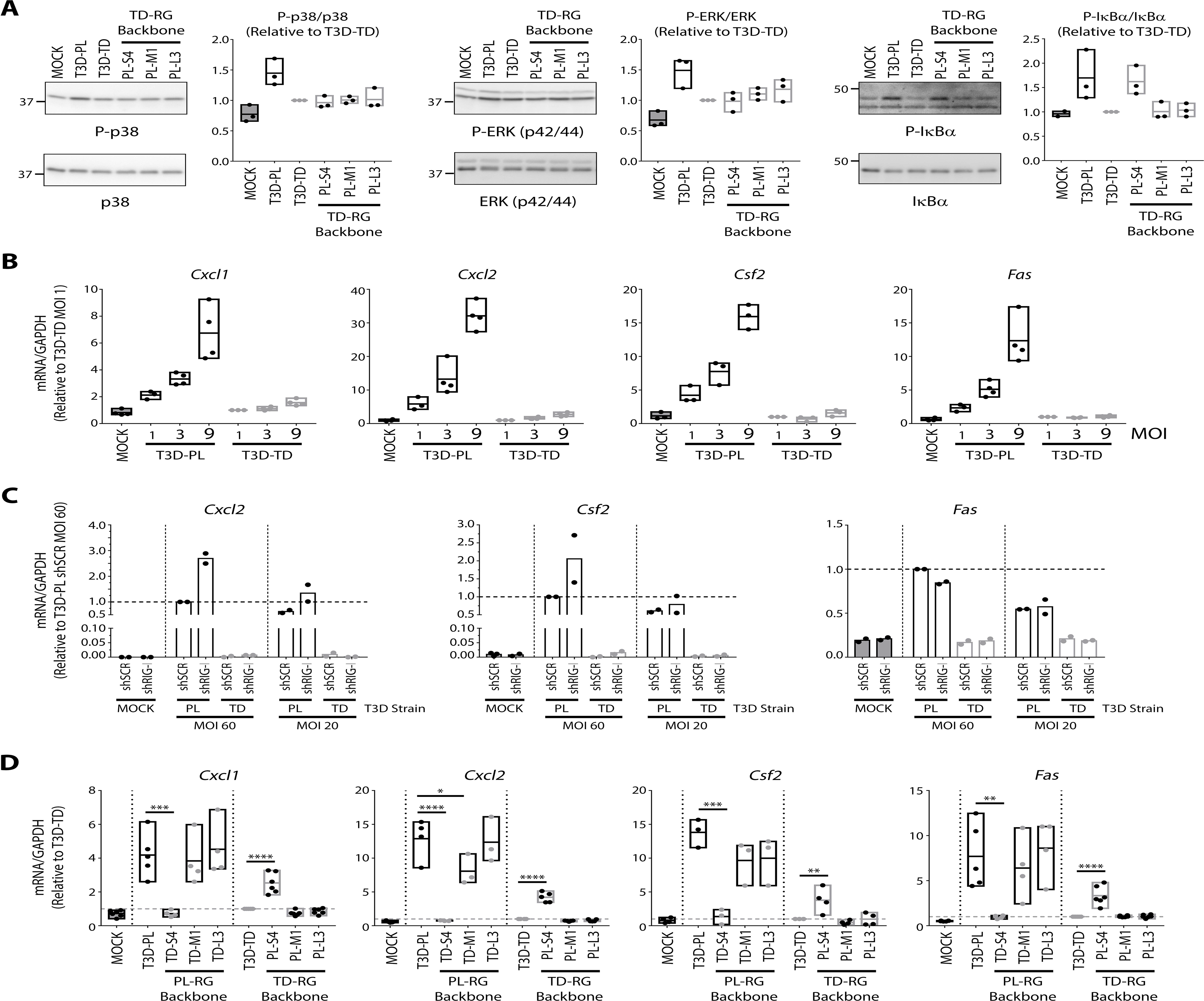
T3D^PL^ S4-encoded σ3 stimulates expression of NFκB-dependent but IFN-independent cytokines. **(A)** Standardized for equal infection (MOI 3), L929 cells were infected for 12hrs with parental T3D^PL^ or T3D^TD^, and S4, M1 and L3 gene monoreassortant TD-RG viruses. (Top) Total cell lysates were collected at 12hpi for Western blot analysis using indicated antibodies to identify reovirus proteins. (Bottom) Densitometric band quantification of phosphorylated relative to total protein. n = 3. **(B)** L929 cells were infected with T3D^PL^ and/or T3D^TD^ at indicated MOI and incubated at 37°C for 12 hours. Total RNA was extracted, converted to cDNA and gene expression relative to housekeeping gene GAPDH was quantified using qRT-PCR. All values were normalized to T3D^TD^ MOI 1. n = 4. **(C)** NIH/3T3 cells stably transduced with scrambled (shSCR) or RIG-I (shRIG) lentivirus were infected with reovirus at indicated MOI (L929 cell line titres) and incubated at 37°C. Samples were collected at 12hpi for RNA extraction, cDNA synthesis and qRT-PCR using gene-specific primers. Values were standardized to corresponding GAPDH and all samples were normalized to shSCR T3D^PL^MOI 60. n = 2. **(D)** Similar experimental outline to (A) except the parental T3D^PL^ or T3D^TD^, and S4, M1 and L3 gene monoreassortant for both PL-RG and TD-RG viruses were assessed. At 12hpi, total RNA was assessed for reovirus S4 RNA expression relative to housekeeping gene GAPDH. n ≥ 3. Statistical significance determined using one-way ANOVA with Dunnett's multiple comparisons test, * p < 0.05, *** p < 0.001, **** p < 0.0001, ns > 0.05.

Serving as transcription factors, NF-κB subunits ultimately stimulate expression of a plethora of NF-κB-dependent genes. The discovery that T3D^PL^ activated RIG-I, IRF3, and IFN-dependent genes less robustly than T3D^TD^ (Figure 5), but reciprocally may activate NF-κB more than T3D^TD^ (Figure 6), raised the possibility that T3D^PL^ (but not T3D^TD^) activated NF-κB-dependent yet RIG-I/IFN-independent genes. This possibility was exciting because NF-κB signaling is typically characterized as being downstream of RIG-I activation, whereas our data potentially introduces a RIG-I independent NF-κB signalling cascade that is differentially stimulated by strains of reovirus with distinct oncolytic potencies. To test this possibility, it was essential that the western blot data be corroborated by data indicating that NF-κB-dependent (and RIG-I independent) genes are indeed stimulated by T3D^PL^ but not T3D^TD^. Microarray and bioinformatics analysis was therefore conducted to identify T3D^PL^- and NF-κB-regulated genes that were RIG-I and IFN-independent. First, we conducted whole genome microarray analysis for NIH3T3 cells that were mock infected, or infected with T3D^PL^ (MOI=60), and focused on genes that were up-regulated by ≥2-fold in T3D^PL^ infected cells relative to mock infection (Supplemental Figure 4). Microarray analysis was also conducted for T3D^PL^-infected NIH3T3 cells stably transduced with shRIG-I, and genes upregulated by reovirus were further subdivided into those whose expression was suppressed by RIG-I knock-down (RIG-I-dependent, cluster 1) versus those that were independent of RIG-I status (RIG-I-independent, cluster 2). To then determine which genes in each cluster are NF-κB-dependent, we made use of a publically available microarray dataset where lipopolysaccharide (LPS) was used to induce both NF-κB and IFN pathways. The database compared LPS-induced gene expression in wild-type MEFs versus MEFs with NF-κB p65/c-Rel subunit knock-out or with IFN receptor (IFNAR) knock-out, to distinguish NF-κB-dependent versus IFN-dependent genes ((Cheng et al., 2017), GEO:GSE35521). Using this public dataset, we further classified T3D^PL^-upregulated genes into four groups: genes that are RIG-I-dependent, NF-κB-dependent, and IFNAR-dependent (cluster 4), RIG-I-dependent, NF-κB-dependent, and IFNAR-independent (cluster 3), RIG-I-independent, NF-κB-dependent, and IFNAR-dependent (cluster 5), and RIG-I-independent, NF-κB-dependent, and IFNAR-independent (cluster 6). Cluster 6 represented T3D^PL^-upregulated genes predicted to be independent of both RIG-I and IFN signalling, but dependent on NF-κB activation, and was therefore chosen for empirical analysis.

Included in cluster 6 were Cxcl1, Csf2, Cxcl2, and Fas. To confirm that these four genes were independent of IFN, L929 cells were treated with IFNα or IFNβ for 12 hours, and gene expression monitored by qRT-PCR (Supplemental Figure 5). As expected, IFN-dependent genes such as Mx1, Cxcl10, Rsad2, Ccl4, Ifi44 and IL6 were upregulated by exposure to IFNs. Conversely, Cxcl1, Csf2, Cxcl2, and Fas genes were not upregulated by IFN treatment, suggesting they are indeed IFN-independent genes as the bioinformatics analysis suggested. When levels of Cxcl1, Csf2, Cxcl2, and Fas were compared between L929 cells infected with T3D laboratory strains at MOIs of 1, 3, and 9 for 12hpi, strong induction (up to 30-fold) was evident in a dose-dependent manner during infection by T3D^PL^ but not T3D^TD^ (Figure 6B). Cxcl1, Csf2, Cxcl2, and Fas induction was also RIG-I independent, since these genes were upregulated by T3D^PL^ to similar extent in NIH3T3 cells transduced with shSCR or shRIG-I (Figure 6C). Moreover, analysis of gene expression among S4, M1, and L3 mono-reassortants showed a clear correlation between expression of these NF-κB-dependent RIG-I/IFN-independent genes and presence of the T3D^PL^-derived S4/σ3 (Figure 6D). Most surprising about the findings, is that NFκB and IRF3 signalling are inversely activated by T3D laboratory strains; T3D^PL^ caused high expression of NFκB-dependent genes and low expression of RIG-I/IFN-dependent genes, but the reciprocal scenario occurred for T3D^TD^. These two signalling pathways are often linked, for example it was recently shown that serotype 3 (T3D) activates both NFκB and IRF3 more than serotype 1 (T1L) (Stuart et al., 2018). While many studies show NFκB and IRF3 downstream of cytosolic sensors like RIG-I, our data suggests an independent source of NFκB signalling that is modulated by σ3. The inverse induction of NFκB – versus RIG-I/IFN-dependent genes by T3D^PL^ versus T3D^TD^ was recapitulated in all four cells lines that we evaluated, including NIH/3T3 (n=2), L929 (n=3), B16-F10 (n=2), and ID8 (n=1), suggesting a widespread phenomenon (data not shown). As will be extrapolated in the discussion, that minor modifications to viral genomes can produce distinct cytokine expression landscapes could be relevant for optimizing virus-induced anti-tumor immunity and for understanding how closely-related viruses cause distinct pathogenic outcomes.

Finally, although NFκB signalling was previously found necessary for induction of cell death by T3D^PL^ (Pan et al., 2011), we determined the effects of modulating NFκB signalling on T3D^PL^ versus T3D^TD^ (Supplemental Figure 5B). While addition of InSolution™ NF-κB Activation Inhibitor decreased cell death following T3D^PL^ infection in a dose-dependent manner, addition of TNF to induce NF-κB activation reciprocally increased cell death following T3D^TD^ infection. Moreover, InSolution™ NF-κB Activation Inhibitor also decreased levels of reovirus proteins (Supplemental Figure 5C), suggesting a role in virus replication in addition to cell death induction. Furthermore, pre-incubation with InSolution™ NF-κB Activation Inhibitor for 3 hours prior to infection reduced virus protein accumulation and cell death by either laboratory strain. Together, the data suggests that both T3D^PL^ and T3D^TD^ benefit from basal activity of NFκB, but that T3D^PL^ can further activate and therefore benefit from NFκB stimulation.

### Specific polymorphisms in S4, M1 and L3 of T3D^PL^ relative to T3D^TD^ have positive or negative effects on replication; and a hybrid between T3D^PL^ and T3D^TD^ has superior oncolytic activity over either T3D laboratory strain

Having identified the genetic and phenotypic contribution of S4, M1, and L3 genes for enhanced oncolysis by T3D^PL^ relative to T3D^TD^, but given that each of these genome segments had 2-3 amino acid differences between the two laboratory strains, we sought to pinpoint the precise amino acids that define oncolytic potency of T3D^PL^. For each polymorphism, the T3D^PL^-derived amino acid was individually cloned into the otherwise T3D^TD^ genetic background. In our nomenclature, the T3D^TD^ amino acid is listed first, then the numerical location of the amino acid, followed by the amino acid identity introduced from T3D^PL^ (e.g., W133R indicates that the T3D^TD^-derived tryptophan was replaced by a T3D^PL^-derived arginine into an otherwise T3D^TD^ genomic background). If the sequence difference indeed contributes to the enhanced oncolytic potency of T3D^PL^, then we expected that the substitution would increase plaque size relative to the T3D^TD^ parental strain.

A surprising result came from segregating the 3 individual variations in the S4-encoded σ3 protein; some amino acid variations increased plaque size, while others decreased plaque size. Among the three amino acid differences at positions 133, 198, and 229, the W133R modification alone was sufficient to almost fully restore plaque size of T3D^TD^ to T3D^PL^ (Figure 7A). Surprisingly, G198K or E229D substitutions marginally reduced plaque size relative to T3D^TD^, and strongly reduced plaque size relative to T3D^TD^ containing the W133R mutation. These experiments therefore indicated that the increased plaque size of T3D^PL^ is attributable to the arginine at position 133 of σ3, while the lysine and aspartic acid at positions 198 and 229 of T3D^PL^ actually suppress plaque size potential.

**Figure 7.**
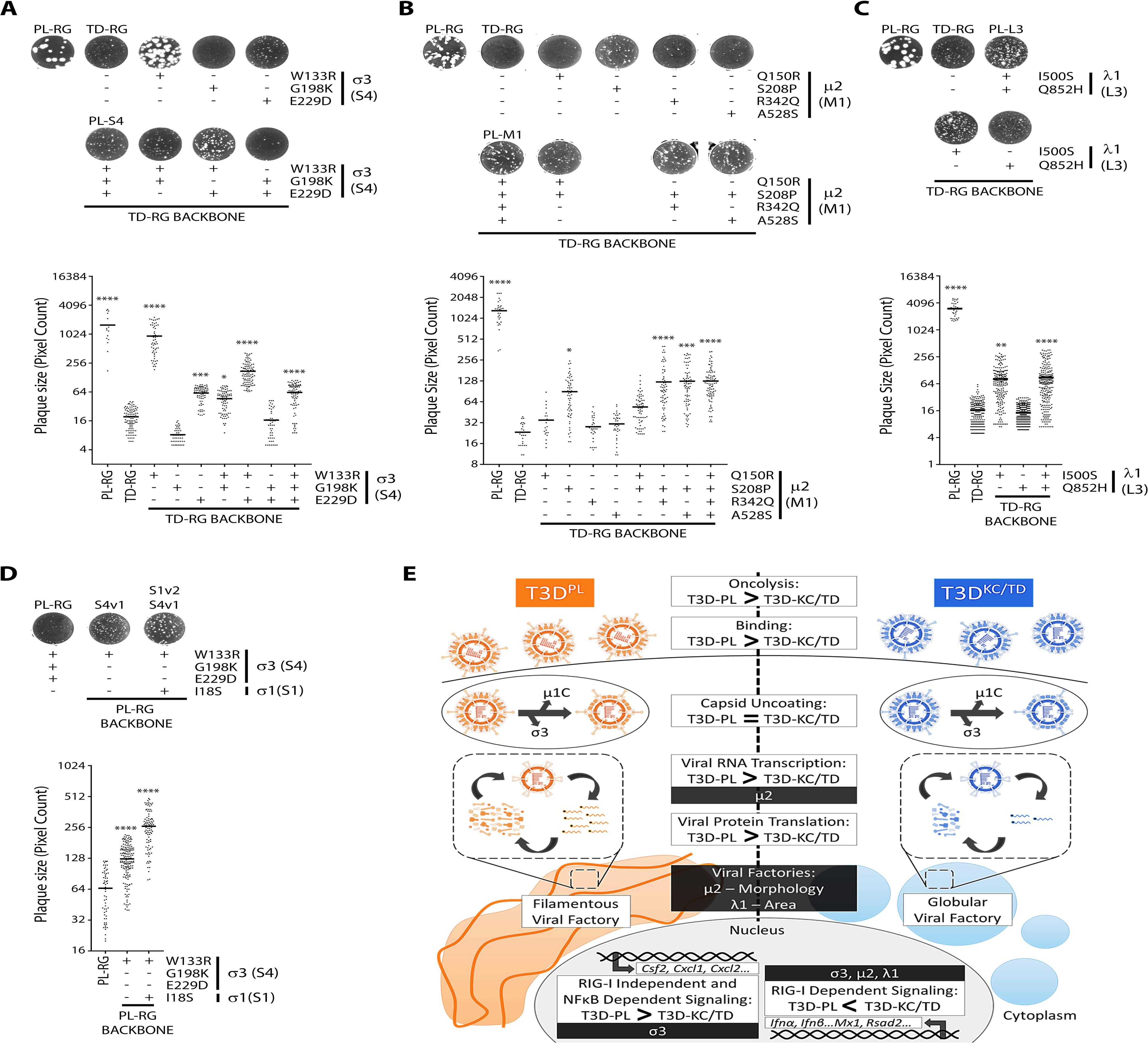
Specific polymorphisms in S4, M1 and L3 of T3D^PL^ relative to T3D^TD^ have positive or negative effects on replication; and a hybrid between T3D^PL^ and T3D^TD^ has superior oncolytic activity over either T3D laboratory strain. **(A-C)** T3D^PL^ polymorphisms were inserted into T3D^TD^ backbone plasmids using site-directed mutagenesis, and modified plasmids were used to generate recombinant viruses in a TD-RG backbone using reverse genetics. Using L929 cells, plaques of parental and recombinant reoviruses were visualized following crystal violet staining and quantified using ImageQuantTL colony counting add-on. (A) S4 gene polymorphisms (B) M1 gene polymorphisms (C) L3 gene polymorphisms. **(D)** Similar experimental outline to (A-C) except specific recombinant viruses were generated in a T3D^PL^-reverse genetics (RG) backbone. Statistical significance determined using one-way ANOVA with Dunnett's multiple comparisons test, * p < 0.05, ** p < 0.01, *** p < 0.001, **** p < 0.0001. In (A), T3D^PL^-RG and S4 (+W133R) were compared to T3D^TD^-RG, whereas remaining samples were independently compared to T3D^RG^. Samples in (B) and (C) compared to T3D^TD^-RG. **(E)** Summary model highlighting genetic and phenotypic differences between T3D^PL^ and T3D^TD^.

The M1-encoded μ2 contains four amino acid differences between T3D^TD^ and T3D^PL^; Q150R, S208P, R342Q, and A528S. The most prominent contributor to plaque size was the S208P polymorphism (Figure 7A). As mentioned, a serine-to-proline difference at position 208 of μ2 was previously described for T1L versus T3D^TD^ and changed the morphology of virus factories (Parker et al., 2002). Q150R, R342Q, and A528S each had marginal contribution to plaque size on their own, but both R342Q, and A528S increased plaque size when combined with S208P relative to S208P alone. Our analysis therefore suggests that in addition to S208P, R342Q and A528S are key contributors to the large plaque phenotype of T3D^PL^ (Figure 7B). Unfortunately, the crystal structure of μ2 has not yet been solved, so it is not possible to map these polymorphisms in tertiary structure. However, both R342Q, and A528S lie near the regions proposed to be involved in NTPase activity and putative nucleotide binding (Eichwald et al., 2017; Kim et al., 2004; Noble and Nibert, 1997b), so these changes could help explain the strongly increased transcriptional activity imparted by the T3D^PL^ M1 gene.

There are two amino acid differences between T3D^TD^ and T3D^PL^ in the L3-encoded λ1 protein; I500S and Q852H. Only the I500S modification increased plaque size in an otherwise T3D^TD^ genomic background, implicating λ1 amino acid 500 as a determinant of enhanced T3D^PL^ replication relative to T3D^TD^ (Figure 7C). On the crystal structure of reovirus core proteins, the I500S mutation maps to the interface between λ1 and σ2. The potential impact of the λ1-σ2 on core assembly and subsequent accumulation of reovirus macromolecules in viral factories, to produce larger factory size will be explored in the discussion.

Finally, our discovery that the σ3 of T3D^PL^ had two variations that were actually restricting T3D^PL^ plaque size, suggested that it may be possible to further increase the oncolytic potency of T3D^PL^ by creating a hybrid sequence between the laboratory strains. Specifically, the data suggested that an arginine at position 133 of σ3 is strongly favorable for in vitro oncolysis relative to a tryptophan, while residues 198 and 229 are most favorable as glycine and glutamic acid from the T3D^TD^ sequence, respectively. If this is true, then we predicted that introduction of glycine at position 198 and glutamic acid at position 229 should further increase plaque size of T3D^PL^. Indeed, plaque size of T3D^PL^ was further increased by K198G and D229E mutations in σ3 (Figure 7D). These findings suggest that this hybrid between T3D^PL^ and T3D^TD^ (T3D^PL/σ3K198G/σ3D229E^) exhibits even higher *in vitro* oncolytic activity than either parental strain, and supports a potential for further improving the activity of T3D^PL^ by genomic manipulation. Lastly, our exploration of σ1 status (Figure 3B-D) suggested that T3D^TD^ had ~50% particles with insufficient σ1 cell attachment proteins for efficient binding (i.e., <3), while T3D^PL^ had a predominance of particles with 7-12 σ1 trimers. Previous studies showed that having 3-7 σ1 per reovirus was optimal for reovirus replication initiation in tumor cells, and could be achieved by a serine-to-isoleucine mutation at position 18 (S18I) of σ1 optimal for efficient entry. The S18I mutation was therefore introduced into T3D^PL/σ3K198G/σ3D229E^, as a strategy to maintain sufficient σ1 for binding but not too many σ1 for efficient entry. The product, T3D^PL/σ3K198G/σ3D229E/ σ1D229E^, exhibited further increase in plaque size (Figure 7D). Altogether our studies demonstrate the genomic and phenotypic contribution of S1, S4, M1, and L3 in reovirus oncolytic potency, and demonstrate that the ideal oncolytic vector may actually be a hybrid between laboratory strains.

## DISCUSSION

Our detailed analysis of reovirus laboratory strains showed that very closely related strains can differ substantially in *in vitro* and *in vivo* oncolytic potency, revealed the mechanisms for enhanced replication in cancer cells associated with superior-oncolytic strains, and discovered unanticipated differences in virus-induced cytokine profiles that could further contribute to the immunotherapeutic aspect of oncolytic reovirus treatment (Figure 7E). These findings provide a cautionary reminder that very small changes in viral proteins can have a very large impact. In recent years, the large market for cancer immunotherapies has provoked a focus on adding immune-stimulating genes to oncolytic viruses (e.g. tumor antigens or GM-CSF), or on combination therapy of oncolytic viruses with checkpoint blockade agents. Our study demonstrates the potential of single amino acid differences alter oncolytic potency, and even to change the cytokine landscape, both of which could affect immunotherapeutic activity. As oncolytic virus studies across the world use independently propagated virus stocks, our findings at minimum propose that the most-oncolytic strain of each virus should be identified and shared for research studies related to cancer therapy; in the case of reovirus, that T3D^PL^ be used as a comparative. Given the increasing ease of sequencing, careful documentation of virus genome sequences would be an important step forward.

Reovirus T3D^PL^ laboratory strain showed substantially better replication than T3D^TD^ on a panel of cancer cell lines that clearly correlated with *in vivo* control of aggressive melanoma (Figure 1A-F). If our goal is to develop the strongest virus oncolytic therapy, then ideally the reovirus T3D^PL^ base vector should continue to undergo development towards utmost oncolytic activity. With respect to improving the oncolytic potency of T3D^PL^, we found that a hybrid between T3D^PL^ and T3D^TD^ laboratory strains (T3D^PL/σ3K198G/σ3D229E/ σ1D229E^) has superior oncolytic activity *in vitro* (Figure 7E). For herpesvirus, it was suggested that “the *in vitro* cytolytic properties of OVs [oncolytic viruses] are poor prognostic indicators of *in vivo* antitumor activity” (Sobol et al., 2011); but it should not be assumed that all viruses are equally maximized. In fact, we previously demonstrated that single amino acid mutations in T3D^PL^, which promote capsid disassembly in tumor cells, can enhance reovirus oncolysis both *in vitro* and *in vivo* (Mohamed et al., 2015; Shmulevitz et al., 2012). Our current data further supports that the reovirus genome has not yet achieved maximal adaptation towards tumors. Keeping in mind that reovirus naturally evolved to be exquisitely well-adapted towards enteric infections, and that the ileum of a gut is quite distinct from a tumor, it seems unsurprising that modifications to reovirus genomes could enhance replication in tumors, dissemination to metastatic sites, and/or induction of anti-tumor immunity. It will be important to test the oncolytic activity of T3D^PL/σ3K198G/σ3D229E/ σ1D229E^ in several animal tumor models, and if it is superior to T3D^PL^ *in vivo*, this variant could function as the next reovirus base vector for further development.

In addition to differences in replication, reovirus laboratory strains induced distinct profiles of virus-stimulated cellular immunomodulatory genes (IMGs). Remarkably, T3D^**TD**^ caused upregulation of IFN-induced IMGs such as CCL5, CCL4, and CXCL10 (Supplementary Figure 3D), while conversely T3D^**PL**^ caused upregulation of IFN-***in***dependent but NF-κB-dependent IMGs such as CXCL2 and CSF2 (Figure 6B, 6C). Reovirus-mediated oncolysis is a combination of both direct virus-induced cytolysis (i.e. demonstrated by tumor clearance in many immunodeficient models (Alain et al., 2002; Hata et al., 2008; Norman et al., 2002a; Sei et al., 2009; Yang et al., 2004) and anti-tumor immune stimulation (Clements et al., 2015; Gujar et al., 2013; Gujar and Lee, 2014; Gujar et al., 2011; Prestwich et al., 2009; Steele et al., 2011). Given that cytokines, chemokines, and innate signalling molecules strongly influence anti-tumor immunity (Adler et al., 2017; Izzi et al., 2014; Romee et al., 2014; Showalter et al., 2017; Waldmann, 2017), we predict that they will impact anti-tumor immunity during reovirus oncolysis. Our data suggests a close investigation into the roles of IFN-dependent versus IFN-independent cytokines on reovirus-strain-specific anti-tumor activity. Based on the 5 cytokines we analyzed, it is impossible to predict the impact of the distinct cytokine profile induced by each strain. As an example, CXCL10 is proposed to have both pro- and anti-proliferative action on Breast cancer depending on the receptors it engages (Liu et al., 2011). Most importantly, many more IFN-dependent and -independent cytokines are likely upregulated differentially by reovirus strains that were not evaluated in this study. A systematic approach will be necessary to determine precisely the benefits or costs of using reovirus strains that induce IFN-dependent versus IFN-independent cytokines. Should both IFN-dependent and -independent IMGs prove important during oncolytic virus treatment, then combination therapy with both T3D^**PL**^ and T3D^**TD**^ reovirus strains (or a hybrid variant that induces both types of signalling) warrants consideration. One must also consider that such cytokines would contribute to heightened clearance of the virus, and therefore to achieve a balance between strong virus replication and strong anti-tumor activity, immunologically distinct oncolytic viruses may need to be sequentially used. Overall, our findings suggest that apart from adding genes to oncolytic viruses to stimulate immunity, small genomic differences among virus strains could impact the immunotherapeutic potential of an oncolytic virus.

Understanding the mechanisms of IFN signalling induction by viruses is important for both virus oncolysis and virus pathogenesis. With respect to IFN-dependent genes, countless examples show that many viruses express viral genes that counteract cellular signalling. Indeed even for reovirus, previous reassortants analysis between reovirus serotypes (T1L) and 3 (T3D) suggested that S2, S4, M1 and L2 reovirus genes function to regulate IFN signalling (Beattie et al., 1995; Bergeron et al., 1998; Sandekian and Lemay, 2015; Sherry, 2009; Sherry et al., 1998; Zurney et al., 2009). But our studies suggest an alternative mechanism affects activation of IFN-dependent IMGs; replication of mono-reassortant and parental viruses was inversely correlated with IFN induction (Figure 5E). We therefore propose an alternative model: T3D^TD^, via delayed onset of replication kinetics, lead to accumulation of viral PAMPs and therefore detection by cellular genes such as RIG-I and MDA5 that activate IFNs, prior to achieving sufficient localized expression of virus proteins that conceal de-novo PAMPS (such as σ3 which binds dsRNA portions of RNAs) or usurp host IRFs. In other words, if incoming reovirus cores fail to establish a virus factory around them quickly, there is less protection from cellular proteins.

In this model, virus replication would inversely correlate with IFN induction. Our findings are very timely, as there is growing interest in the concept that defective virions in a virus quasispecies contribute to innate detection of viruses, since such defective virions are debilitated in their ability to dampen the immune response (reviewed in (Lopez, 2014, 2019)). If our model is correct, then similar to T3D^TD^, other virus variants in a quasispecies with slow kinetics of establishing infection may be inherently more vulnerable to innate detection, which could help protect from rapid-replicating variants in the quasispecies, but may also increase immune-associated pathogenesis; for example in Ebola and Zika, for which pathogenesis correlates with heighted innate signalling response (Ebihara et al., 2006; Tripathi et al., 2017).

With respect to IFN induction, it seemed also paradoxical that co-infections with T3D^PL^ and T3D^TD^ still produced high IFN signalling (i.e. phenocopy of T3D^TD^ with respect to IFN activation), yet high replication of virus (i.e. phenocopy of T3D^PL^ with respect to virus macromolecular synthesis), suggesting it unlikely that T3D^PL^ globally counteracts the IFN signalling. Reovirus incoming cores were previously demonstrated to establish virus factories around them (Broering et al., 2004; Broering et al., 2000a), so during co-infections, we predict that incoming T3D^PL^ and T3D^TD^ cores each establish their own factories. In this scenario, the slow-replicating T3D^TD^ virus would still cause induction of the IFN signalling pathway. But since IFN signalling does not affect the first round of reovirus replication (Figure 5F-H, Supplementary Figure 3B-D), induction of IFN by T3D^TD^ would not impact the replication of T3D^PL^, thus explaining the continued high levels of viral proteins during T3D^PL^/T3D^TD^ co-infections. T3D^TD^-mediated IFN induction would presumably affect cell-cell spread of either T3D^PL^ or T3D^TD^, since IFN was inhibitory to both strains through paracrine fashion (Figure 5H). The results beckon a better understanding of how incoming cores establish factories, and whether individual incoming variants have distinct consequences in a co-infected cell.

Our phenotypic comparison of the highly oncolytic T3D^PL^ to the less-oncolytic T3D^TD^ also revealed four virus attributes that yield a strong oncolytic reovirus base vector (Figure 7E). First, T3D^PL^ exhibited higher binding to tumorigenic cells than T3D^TD^ (Figure 3D). Cell attachment is mediated by reovirus outer capsid σ1 protein (Lee et al., 1981). We and others previously demonstrated that reovirus particles can house from 0 to 12 σ1 molecules, and that 3 or more σ1 molecules are sufficient for binding to cells (Larson et al., 1994; Mohamed et al., 2015). We now show that T3D^TD^ has higher proportion of particles with two or less σ1 molecules, which explains the reduced cell binding capacity of this strain. The difference in binding between the two strains was reproducible but small, and therefore not surprisingly the genes previously shown to affect σ1 level (i.e. σ1 or λ2 proteins that anchor σ1) did not robustly assort with high oncolytic activity (Figure 2B, 2D, 2E). Second, and most strikingly, T3D^PL^ established rapid onset of viral RNA and protein synthesis, leading to more rapid kinetics of progeny virus production and a larger virus burst size (Figure 3E-H, Supplementary Figure 2). This points to the importance of rapid onset of infection for oncolysis. Thirdly, T3D^PL^ also establishes globular factories, and while the importance of globular over filamentous factories remains to be fully characterized, it was clear that variations in L3-encoded λ1 increase the size of globular factories likely reflecting more-efficient virus amplification (Figure 4H). Fourthly, T3D^PL^ more-rapidly caused cell death and efficiently disseminated (Figure 3E, 3I, 3J, Supplementary Figure 2). We also noted strong differences in antiviral signaling between T3D^PL^ and T3D^TD^ as described above. Put together, our findings suggest that with respect to direct oncolysis, an oncolytic reovirus vector should have these attributes. In future, mono-reassortants should be compared *in vivo* to determine which of the virus attributes contribute most to *in vivo* oncolysis; though we predict that the mono-reassortants will show intermediate phenotypes between T3D^PL^ and T3D^TD^ parental strains as they did for plaque size.

Our genetic analysis has uncovered three reovirus genes that prominently contribute to the enhanced oncolytic activity of T3D^PL^ in vitro: M1 (encoding μ2), L3 (encoding λ1), and S4 (encoding σ3). Functional studies then revealed several phenotypic properties controlled by these proteins that likely contribute to the enhanced potency of T3D^PL^. μ2 enhanced the transcriptional activity of viral cores and influenced viral factory morphology, λ1 increased factory size, and σ3 was required for induction of NF-κb-dependent, IFN-independent genes. A genotypic comparison was then used to identify reovirus genes that strongly associated with the high oncolytic activity of T3D^PL^ relative to T3D^TD^. First, the M1-encoded μ2 protein of T3D^PL^ was demonstrated to increase virus transcription activity and promote a globular morphology of reovirus factories (Figure 4D, 4F-H). The μ2 protein serves at least two distinct functions; a structural protein within virus cores that functions as a transcription co-factor, and a tubulin-associated protein during virus replication involved in accumulating virus proteins into localized sites of virus replication called factories (Coombs, 1998; Parker et al., 2002). This is the first time, to our knowledge, that a mutation in μ2 is shown to increase rates of transcription in infected cells, and supports previous suggestions that μ2 is a polymerase co-factor (Kim et al., 2004; Noble and Nibert, 1997b). Previous studies also proposed that μ2has NTPase activity and amino acids in positions 399-421 and 446-449 affect nucleotide binding and NTPase activities (Eichwald et al., 2017; Kim et al., 2004; Noble and Nibert, 1997b). We found that the S208P mutation is most-important for increased replication of T3D^PL^, but that the other mutations (R342Q and A528S) provide especially when combined with the S208P mutation (Figure 7B). None of these variations lie in the so-far identified NTP binding and NTPase activity domains, suggesting that either the mutations affect these activities in the folded μ2 structure, or that additional domains impact the transcriptional co-factor role of μ2. Future crystal structure determination of μ2 would help decipher between these possibilities. With respect to our finding that all reassortants containing T3D^PL^ μ2 exhibited globular factory morphology, this phenotype was already previously described to be associated with polymorphisms at position 208 (Parker et al., 2002; Yin et al., 2004). Specifically, other laboratories noticed that a proline at position 208 of μ 2 produced globular factory morphology for reovirus serotype 1 (T1L), while a serine at the same position produced fibrous factory morphology for reovirus serotype 3 (T3D). Since T3D^PL^ has a proline at 208 and globular morphology, we predict that the T3D strain used in previous analyses was of T3D^TD^ origin. Altogether, our findings suggest that strong transcription activity and globular factory morphology associate with high oncolytic potency. While it is clear how enhanced transcription would permit rapid onset of virus infection and therefore oncolysis, additional studies are needed to understand how globular-versus-fibrous factory morphology could promote reovirus oncolytic potency.

A second gene prominently attributed to stronger oncolytic activity of T3D^PL^ was λ1. The inner capsid of reovirus cores is composed of dimers of λ1 and σ2 (Supplemental Figure 6). T3D^PL^ and T3D^TD^ have two amino acid differences in λ1 (I500S and Q852H), but only the I500S polymorphism corresponded with increased reovirus replication in tumor cells (Figure 7C). The λ1 gene of T3D^PL^ was also associated with strikingly larger virus factories, without affecting the morphology of the virus factory which depended on μ2 as discussed above. Based on crystal structures of reovirus cores (Reinisch et al., 2000), amino acid 500 is located on the surface of λ1 that associated with σ2. The most-likely hypothesis is therefore that I500S promotes λ1-σ2 interactions and thereby co-association of these core proteins to factories and assembly of new cores, which would serve to amplify reovirus replication by synthesizing more RNAs. Alternative possibilities are that λ1 I500S mutation contributes to virus protein trafficking to factories or factory construction.

The third reovirus gene that correlated with superior oncolysis by T3D^PL^ was σ3. Analysis of the specific polymorphisms of σ3 between T3D^TD^ and T3D^PL^ led to a surprising result; while T3D^PL^ has one polymorphism that drastically promotes reovirus replication (W133R), the other two polymorphisms (G198K and E229D) are unfavorable and reduce reovirus replication relative to the W133R polymorphism alone (Figure 7A). As discussed above, this finding proposes that the optimal reovirus base vector might consist of a T3DPL/TD hybrid, whereby all sequences are of T3D^PL^ origin except σ3 which would contain a glycine at 198 and glutamic acid at 229 derived from T3D^TD^. The σ3 protein has two well-characterized roles: it is the outer-most capsid protein that maintains reovirus stability and must be removed during virus entry, and it binds dsRNA during virus replication to quench cellular dsRNA binding proteins such as RNA-dependent protein kinase (PKR) and subdues antiviral mechanisms (Beattie et al., 1995; Chandran et al., 1999; Jane-Valbuena et al., 2002; Olland et al., 2001; Yue and Shatkin, 1997). Interestingly, all three polymorphisms lie in similar proximity in the σ3 structure (Supplementary Figure 6). The polymorphisms are next to the hypercleavable sequence needed for degradation of σ3 during virus entry (Liemann et al., 2002; Mendez et al., 2003), but also among the charged surface of σ3 involved in dsRNA binding (Denzler and Jacobs, 1994; Mabrouk et al., 1995; Mabrouk and Lemay, 1994; Miller and Samuel, 1992; Olland et al., 2001; Schiff et al., 1988; Wang et al., 1996). In our studies, minimal differences were noted in σ3 degradation during entry, but σ3 of T3D^PL^ was clearly associated with induction of NF-κB-dependent, IFN-*in*dependent IMGs (Figure 6A, 6D). We therefore propose that the polymorphisms likely affect reovirus replication by modulating dsRNA-specific but IFN-independent cellular processes. For example, PKR was previously suggested to promote reovirus replication by terminating host translation outside of viral factories, leaving ribosomes to freely translate reovirus mRNAs at virus factories where host anti-viral factors were excluded (Desmet et al., 2014; Smith et al., 2006; Stanifer et al., 2017; Yue and Shatkin, 1997). We propose a model whereby T3D^TD^ predominantly stimulates RIG-I and IRF3 signalling, with minimal NF-κB activation, leading primarily to expression of IFNs and IFN-dependent genes (Figure 7E). Conversely, T3D^PL^ activates another pathway such as PKR that stimulates strong NF-κB activity, but overcomes IRF3 activation through rapid onset of infection, thereby chiefly stimulating expression of NF-κB-dependent genes. As PKR and NF-κB have been previously associated with promotion of reovirus, for example by stimulating host protein shut-off, virus replication and apoptosis of infected cells (Hansberger et al., 2007; Knowlton et al., 2012; Pan et al., 2013; Pan et al., 2011; Smith et al., 2006; Yue and Shatkin, 1997), this model would favor T3D^PL^. Future assessment of σ3 – dsRNA binding efficiencies and activities of additional signalling molecules such upstream of NF-kB, as PKR, will help resolve precisely how the three polymorphisms of σ3 contribute to enhanced IFN-independent gene expression and replication of T3D^PL^.

Altogether this study provides the first comprehensive analysis of how strains of an oncolytic virus can affect oncolytic activity. We show that single amino acid changes drastically affect virus replication and therefore direct viral oncolysis. Increased virus replication and dissemination could also increase opportunities for the virus to stimulate anti-tumor immune responses. Moreover, we show that single amino acid changes also change the profile of immunomodulatory host genes expressed by infected cells, which could also affect recruitment of immune cells important for anti-tumor response. The data supports that strong attention should be paid to the genome and phenotypic characteristics of specific oncolytic virus strains, if to achieve best therapeutic value of oncolytic viruses.

## MATERIALS AND METHODS

### Cell lines

All cell lines were grown at 37°C at 5% CO2 and all media was supplemented with 1x antibiotic antimycotics (A5955, Millipore Sigma). Except for NIH/3T3 media that was supplemented with 10% NCS (N4637, Millipore Sigma), all other media was supplemented with 10% FBS (F1051, Millipore Sigma). L929, NIH/3T3, H1299, ID8, HCT 116 and B16-F10 cell lines (Dr. Patrick Lee, Dalhousie University), Huh7.5 (Dr. Michael Houghton, University of Alberta), WT MEFs, RIG-I/MDA5 -/- MEFs (Dr. Michael Gale Jr, University of Washington) and BHK-21-BSR T7/5 (Dr. Ursula Buchholz, NIAID) were generous gifts. L929 cell line was cultured in MEM (M4655, Millipore Sigma) supplemented with 1× non-essential amino acids (M7145, Millipore Sigma) and 1mM sodium pyruvate (S8636, Millipore Sigma). L929 cell line in suspension were cultured in Joklik’s modified MEM (pH 7.2) (M0518, Millipore Sigma) supplemented with 2g/L sodium bicarbonate (BP328, Fisher Scientific), 1.2g/L HEPES (BP310, Fisher Scientific), 1× non-essential amino acids (M7145, Millipore Sigma) and 1mM sodium pyruvate (S8636, Millipore Sigma). H1299, ID8 and T-47D cell lines were cultured in RPMI (R8758, Millipore Sigma). NIH/3T3, B16-F10, Huh7.5, WT MEFs, RIG-I/MDA5 -/- MEFs and BHK-21-BSR T7/5 cell lines were cultured in DMEM (D5796, Millipore Sigma) supplemented with 1mM sodium pyruvate (S8636, Millipore Sigma). BHK-21-BSR T7/5 cell line was selected in media containing 1mg/ml G418 (A1720, Millipore Sigma) every second passage. All cell lines were routinely assessed for mycoplasma contamination using 0.5μg/ml Hoechst 33352 (H1399, ThermoFisher Scientific) staining or PCR (G238, ABM).

### Reovirus stocks

Seed stock lysates of T1L, T2J, T3D-PL (Dr. Patrick Lee, Dalhousie University), T3D-KC (Dr. Kevin Coombs, University of Manitoba) and T3D-TD (Dr. Terence Dermody, University of Pittsburgh) were gifts in kind. T3D-ATCC (ATCC® VR-824) was purchased from American Type Culture Collection. Reovirus lysates were plaque purified and second or third passage L929 cell lysates were used as spinner culture inoculums.

### Plaque Purification

Media was removed from confluent L929 cell monolayers in 6 well plates. Reovirus lysates were diluted in MEM (no additives) and 200μl of the dilution was added to each well. Virus inoculum was incubated for 1hr at 37°C, while rocking gently every 5-10min. Virus inoculum was removed prior to addition of 3ml agar overlay which constituted an equal ratio of 2% Agar and 2X MEM (Temin's modification, no phenol red, 11935046, ThermoFisher Scientific) supplemented with 20% FBS, 2× non-essential amino acids (M7145, Millipore Sigma), 2mM sodium pyruvate (S8636, Millipore Sigma) and 2x antibiotic antimycotics (A5955, Millipore Sigma). The agar overlays were allowed to solidify for 30 minutes at room temperature and incubated at 37°C until plaques were clearly visible. Isolated plaques were selected and marked on the plate base using a sharpie. Using sterile cotton plugged glass pasteur pipets and rubber bulbs, agar plugs over selected plaques were carefully extracted, dispensed into 1.5ml microcentrifuge tubes with 200μl of MEM (no additives) and incubated overnight at 4°C.

### Reovirus Passaging

Following overnight incubation, plaque isolated agar plug in MEM was diluted in 1:1 in 100μl MEM (no additives). Media was removed from 60-80% confluent L929 cell monolayers in 6 well plates and 200μl of diluted plaque isolated agar plug was added to the well. Following incubation for 1hr at 37°C (rocking gently every 5-10min), 2ml cell culture media was added to each well. Cell death was monitored microscopically every 12-24hrs. When more than 80% cell death was attained, each well was scraped using a sterile plastic cell lifter (08100240, Fisher Scientific) and transferred to 2ml microcentrifuge tube. This was referred to as passage 1. Following 3 freeze/vortex/thaw cycles, virus concentration in passage 1 lysate was determined using plaque assay.

Lysate from passage 1 was serially diluted in MEM (no additives) such that a final dilution of MOI 1-3 (assuming 1.1×107 cells) in 1ml was attained. Media was removed from 60-80% confluent (1.1×107 cells at 80%) L929 cells in a 55cm2 dish and 1ml passage 1 lysate dilution was added. Following incubation for 1hr at 37°C, while rocking gently every 5-10min, 9ml cell culture media was added. Cell death was monitored microscopically every 12-24hrs. When more than 80% cell death was attained, dish was scraped using a sterile plastic cell lifter (08100240, Fisher Scientific) and transferred to a 15ml conical centrifuge tube. This was referred to as passage 2. Following 3 freeze/vortex/thaw cycles, virus concentration in passage 2 lysate was determined using plaque assay.

T3D-PL, T3D-KC and T3D-TD lysates from passage 2 or PL-RG, TD-RG, TD+PL-S4, TD+PL-M1, TD+PL-L3, PL+TD-S4, PL+TD-M1 and PL+TD-L3 lysates from passage 1, were serially diluted in MEM (no additives) such that a final dilution of MOI 1-3 (assuming 3×107 cells) in 6ml was attained. Media was removed from 2 150cm2 dishes with 60-80% confluent (3×107 cells at 80%) L929 cells and 3ml passage 2 lysate dilution was added to each dish. Following incubation for 1hr at 37°C, while rocking gently every 5-10min, 15ml cell culture media was added to each dish. Cell death was monitored microscopically every 12-24hrs. When more than 80% cell death was attained, both dishes were scraped using a sterile plastic cell lifter (08100240, Fisher Scientific), pooled and transferred to a 50ml conical centrifuge tube. Following 3 freeze/vortex/thaw cycles, virus concentration was determined using plaque assay. This passage was used as the inoculum for large scale spinner culture amplification.

### Large Scale Reovirus Amplification

Four 150cm2 dishes with 100% confluent L929 cell monolayers were detached using trypsin, were resuspended in 500ml JMEM culture media and added to sterile 2L flat bottom flask (10-035G, Fisher Scientific) with stir bar and aluminum foil cover. Flask containing cells was incubated at 37°C with low speed stirring in either a large chamber incubator or a water bath, neither of which had CO2 regulation. Cell density was monitored every 24hrs by removing a 1ml aliquot and performing a trypan blue cell count using either a haemocytometer or Biorad TC-20 cell counter. When cell density reached approximately 1×106 cells/ml, 700ml JMEM culture media was added to the flask. Cell density was monitored every 12-24hrs and at approximately 1×106 cells/ml (1.2×109 cells/1200ml/flask), virus lysates were diluted in MEM (no additives) and added to the flask at an MOI 1-3. Cell density and death was monitored every 12-24hrs. At 50-70% cell death, cells were collected by centrifugation at 1000g for 15min at 4°C. The cell pellet from each flask was resuspended in 15ml homogenization buffer (10mM Tris pH 7.4, 250mM NaCl, 10mM β-mercaptoethanol, 1× protease inhibitor cocktail), transferred to a 50ml conical centrifuge tube and frozen at −80°C. Frozen cell pellets were thawed at 4°C and vortexed for 5min on maximum speed. Per cell pellet, 10ml vertel XF (Dymar Chemicals Limited, ON, Canada) was added, vortexed for 5min on maximum speed and probe sonicated for 30sec on maximum amplitude ensuring a homogenous emulsification. Following centrifugation at 4000rpm for 10min at 4°C, the top (reddish-pink color) layer was separated into a new 50ml conical centrifuge tube. Ten ml of vertel XF (Dymar Chemicals Limited, ON, Canada) was added to the top layer, vortexed for 5min on maximum speed and probe sonicated for 30sec on maximum amplitude ensuring a homogenous emulsification. Following centrifugation at 4000rpm for 10min at 4°C, the top (reddish-pink color) layer was separated into a new 50ml conical centrifuge tube and layered on a CsCl gradient (9ml 1.4g/cc, 9ml 1.2g/cc) in SW28 ultra clear tubes. Tubes were topped up with PBS, weight balanced on a digital scale and centrifuged at 100,000g at 4°C for 6-12hrs. Reovirus separates into two white bands signifying top component empty virions and bottom layer full virions. The bottom band was extracted using a needle (19-22Ga) and syringe (5-10ml) and loaded into dialysis tubing (10-20KDa). Extracted virus was dialyzed in dialysis buffer (150mM NaCl, 10mM MgCl2, 10mM Tris pH 7.4) with 3 buffer changes, the first after 1-3hrs, the second after 3-6hrs and third after 6-12hrs. The dialyzed virus was transferred into a 15ml conical centrifuge tube and allowed any precipitate to settle at 4°C for 12hrs. The supernatant was aliquoted into 1.5ml microcentrifuge tubes and stored at 4°C. Virus concentration was determined using plaque assay

### Reovirus plaque assays

Reovirus dilutions were added to 100% confluent L929 cells for 1hr at 37°C with gentle rocking every 10 minutes, followed by addition of agar overlay (1:1 ratio of 2% agar 2× JMEM media). Overlays were allowed to solidify for 30 minutes at room temperature and incubated at 37°C. When plaques became visible (3-7 days post infection), 4% formaldehyde solution (33314, Alfa Aesar) was added to the overlay for 30 minutes. Formaldehyde was discarded, agar overlays were carefully scooped out and cells were further fixed with methanol for 5min. Methanol was discarded, and cells were stained with crystal violet solution (1% crystal violet (C581, Fisher Scientific) in 50% ethanol and 50% water) for 10min and rinsed with water. For cell lines other than L929, after methanol staining, plaques were stained using immunocytochemistry with rabbit anti-reovirus pAb. Plaques were scanned on the ImageQuant LAS4010 imager (GE Healthcare Life Sciences), and plaque area was measured using ImageQuant TL software (GE Healthcare Life Sciences).

### Virus Infections

Reovirus (purified stock, cell lysate, plaque isolated agar plug) was diluted in MEM (no additives). Cell culture media was aspirated from 90-100% confluent cell monolayers. Virus inoculum was added to each well and incubated for 1hr with gentle rocking every 5-10min to allow for virus adsorption. The virus inoculum was aspirated and the monolayer was washed once with MEM (no additives). Cell culture media was added and the incubated at 37°C. Note that a “wash” in the above protocol constitutes addition of reagent, gently rocking back and forth twice and aspiration of reagent.

When samples were assessed and or standardized for virus binding, cells (90-100% confluent cell monolayers) were pre-incubated at 4°C for 30min. Cell culture media was aspirated, virus inoculum was added to each well and incubated at 4°C for 1hr with gentle rocking every 5-10min to allow for virus adsorption. The virus inoculum was aspirated and the monolayer was washed twice with ice-cold MEM (no additives). Cell culture media was added and the incubated at 37°C. During virus adsorption, the cell plates, virus inoculum and MEM (no additives) were always kept on ice or the 4°C fridge. Note that a “wash” in the above protocol constitutes addition of reagent, gently rocking back and forth twice and aspiration of reagent.

### Immunocytochemistry

Colorimetric infectivity assay: Following reovirus infection (Virus infection or Reovirus plaque assay protocols), cell culture media was aspirated, or agar overlays were removed, and cell monolayers were washed once with PBS. Cells were fixed with methanol or 4% paraformaldehyde, washed twice with PBS, and incubated with blocking buffer (3% BSA/PBS/0.1% Triton X-100) for 1 hour at room temperature. Blocking buffer was aspirated and primary antibody (rabbit anti-reovirus pAb) diluted in blocking buffer for a final concentration of 1:10,000 was added and incubated overnight at 4°C. Samples were washed 3 × 5minutes with PBS/0.1% Triton X-100. Secondary antibody diluted in blocking buffer (goat anti-rabbit AP) was added and incubated for 1-3hrs at room temperature. Samples were washed 3 × 5minutes with PBS/0.1% Triton X-100. NBT/BCIP substrate diluted in AP buffer (100mM Tris pH 9.5, 100mM NaCl, 5mM MgCl2) was added and infected cells were monitored for black/purple staining using microscopy. 100× NBT/BCIP substrate stocks were diluted as follows in dimethylformamide (DMF) (D4551, Millipore Sigma): NBT (30mg/1ml) (B8503, Millipore Sigma), BCIP (15mg/1ml) (N6639, Millipore Sigma). When infected cells had stained a dark purple color, substrate was removed and reactions were stopped by adding PBS/5mM EDTA. Note that a “wash” in the above protocol constitutes addition of reagent, gently rocking back and forth twice and aspiration of reagent.

### Fluorescence infectivity assay

Following reovirus infection (Virus infection protocol), cell culture media was aspirated, and cell monolayers were washed once with PBS. Cells were fixed with 4% paraformaldehyde, washed twice with PBS, and incubated with blocking buffer (3% BSA/PBS/0.1% Triton X-100) for 1 hour at room temperature. Blocking buffer was aspirated and primary antibody (rabbit anti-reovirus pAb) diluted in blocking buffer for a final concentration of 1:10,000 was added and incubated overnight at 4°C. Samples were washed 3 × 5minutes with PBS/0.1% Triton X-100. Secondary antibody diluted in blocking buffer (goat anti-rabbit Alexa Fluor 488) was added and incubated for 1-3hrs at room temperature. Samples were washed 3 × 5minutes with PBS/0.1% Triton X-100. Nuclei were stained with 0.5μg/ml Hoechst 33352 (H1399, ThermoFisher Scientific) for 15min and stained samples were visualized and imaged using EVOS FL Auto Cell Imaging System (ThermoFisher Scientific). Note that a “wash” in the above protocol constitutes addition of reagent, gently rocking back and forth twice and aspiration of reagent.

### Immunofluorescence

Cells were seeded on #1.5 thickness coverslips. Following reovirus infection (Virus infection protocol), cell culture media was aspirated, and cell monolayers were washed once with PBS. Cells were fixed with 4% paraformaldehyde, washed twice with PBS, and incubated with blocking buffer (3% BSA/PBS/0.1% Triton X-100) for 1 hour at room temperature. Blocking buffer was aspirated and primary antibody (mouse anti-σNS 3E10 conjugated to Alexa Fluor 568, mouse-anti-σ3 10G10 conjugated to Alexa Fluor 647, rabbit anti-μ2 pAb) diluted in blocking buffer was added and incubated overnight at 4°C. Primary mAbs mouse anti-σNS and mouse-anti-σ3 10G10 were conjugated to Alexa Fluor 568 and Alexa Fluor 647, respectively, using APEX antibody labeling kits as per manufacturer’s protocol (ThermoFisher Scientific). Samples were washed 3 × 5minutes with PBS/0.1% Triton X-100. Secondary antibody diluted in blocking buffer (goat anti-rabbit Alexa Fluor 488) was added and incubated for 1-3hrs at room temperature. Samples were washed 3 × 5minutes with PBS/0.1% Triton X-100. Nuclei were stained with 0.5μg/ml Hoechst 33352 (H1399, ThermoFisher Scientific) for 15min. Coverslips were mounted on microscope slides using 10μl SlowFade Diamond (S36967, ThermoFisher Scientific) and visualized using an Olympus IX-81 spinning disk confocal microscope (Quorum Technologies). Note that a “wash” in the above protocol constitutes addition of reagent, gently rocking back and forth twice and aspiration of reagent.

### Flow cytometry

Cell culture media was aspirated and cell monolayers were rinsed with PBS (1ml/12well). PBS was discarded and trypsin (200μl/12well) was added and incubated until cells detached. Tryspin in detached cells was quenched with cell culture media (1ml/12well). Volumes of PBS, trypsin and cell culture media were scaled up/down according to well size. Detached cells were centrifuged and washed with 1ml PBS. Refer to Table 1.1 for reagent volumes. PBS was aspirated, cell pellet was gently resuspended in 4% paraformaldehyde and incubated at 4°C for 30min. Samples were centrifuged and washed with 1ml PBS/0.1% Triton X-100 (wash buffer). Wash buffer was aspirated, cell pellet was gently resuspended in 3% BSA/wash buffer (blocking buffer) and incubated at 1 hour at room temperature for 30min. Samples were spiked with primary antibody (rabbit anti-reovirus pAb) diluted in blocking buffer for a final concentration of 1:10,000 and incubated overnight at 4°C. Samples were centrifuged and washed twice with 1ml wash buffer. Wash buffer was aspirated, and cell pellet was gently resuspended in secondary antibody diluted in blocking buffer (goat anti-rabbit Alexa Fluor 647 at 1:1:2,000 dilution) and incubated for 1-3hrs at room temperature. Samples were centrifuged and washed twice with 1ml wash buffer. Wash buffer was aspirated, and cell pellet was gently resuspended in 500μl PBS. Samples were processed using a FACSCanto (BD Biosciences) and data was analyzed using FSC Express 5 (De Novo Software). Total cells were gated using FSC-A and SSC-A, while single cells were gated using FSC-A and FSC-H. A minimum of 10,000 total cells were collected for each sample. Note that a “wash” in the above protocol constitutes aspiration of supernatant, resuspension of pellet and centrifugation. Prior to fixation, cells were centrifuged at 500g for 5min at 4°C. After fixation, cells were centrifuged at 1,000g for 5min at 4°C.

### RNA extraction and RT-PCR

Cells were lysed in TRI Reagent® (T9424, Millipore Sigma) and the aqueous phase was separated following chloroform extraction as per TRI Reagent® protocol. Ethanol was mixed with the aqueous phase and RNA isolation protocol was continued as per GenElute Mammalian Total RNA Miniprep kit (RTN350, Millipore Sigma) protocol. RNA was eluted using RNAse free water and total RNA was quantified using Biodrop DUO (Biodrop). Using 1μg RNA per 20μl reaction, cDNA synthesis was performed with random primers (48190011, ThermoFisher Scientific) and M-MLV reverse transcriptase (28025013, ThermoFisher Scientific) as per M-MLV reverse transcriptase protocol. Following a 1/8 cDNA dilution, RT-PCR reactions were executed as per SsoFast EvaGreen Supermix (1725204, Bio-Rad) protocol using a CFX96 system (Bio-Rad).

### Western blot analysis

Cells were washed with PBS and lysed in RIPA buffer (50mM Tris pH 7.4, 150mM NaCl, 1% IGEPAL CA-630 (NP-40), 0.5% sodium deoxycholate) supplemented with protease inhibitor cocktail (11873580001, Roche) and phosphatase inhibitors (1mM sodium orthovanadate, 10mM β-glycerophosphate, 50mM sodium fluoride). For each 12 well, 100μl lysis buffer was used and volume was scaled up/down according to well size. Following addition of 5× protein sample buffer (250mM Tris pH 6.8, 5% SDS, 45% glycerol, 9% β-mercaptoethanol, 0.01% bromophenol blue) for a final 1× protein sample buffer, samples were heated for 5min at 100°C and loaded onto SDS-acrylamide gels. After SDS-PAGE, separated proteins were transferred onto nitrocellulose membranes using the Trans-Blot® Turbo™ Transfer System (Bio-Rad). Membranes were blocked with 3% BSA/TBS-T (blocking buffer) and incubated with primary and secondary antibodies (5ml blocking buffer, primary and secondary antibody per mini (7×8.5cm) membrane). Membranes were washed 3×5min with TBS-T after primary and secondary antibodies. (10ml wash buffer per mini (7×8.5cm) membrane). Membranes with HRP-conjugated antibodies were exposed to ECL Plus Western Blotting Substrate (32132, ThermoFisher Scientific) (2ml substrate per mini (7×8.5cm) membrane) for 2min at room temperature. Prior to visualization, membranes were rinsed with TBT. Membranes were visualized using ImageQuant LAS4010 imager (GE Healthcare Life Sciences), and densitometric analysis was performed by using ImageQuant TL software (GE Healthcare Life Sciences). Note that a “wash” in the above protocol constitutes addition of reagent, gently rocking back and forth twice and removal of reagent.

### Double-stranded genomic reovirus RNA visualization

RNA was extracted from CsCl purified reovius preparations (~1×1010 virus particles) using TRI Reagent LS (T3934, Millipore Sigma) as per manufacturer’s protocol. Purified RNA was diluted in 4× Laemmli sample buffer (1610747, Bio-Rad), and separated on an 8% SDS-acrylamide gel for 22 hours at 6mA (per gel) at 4°C. RNA was stained using ethidium bromide (0.5 μg/mL) for 1hr at room temperature, gels were destained with TAE (40mM Tris, 5 mM sodium acetate, 1 mM EDTA [pH 7.5]) for 2hrs at room temperature and gels were imaged on ImageQuant LAS4010 imager (GE Healthcare Life Sciences).

### Flow cytometry binding assay

L929 cells were detached with CellStripper (Corning), diluted and aliquoted (5×10^10^ cells/sample/ml), pre-chilled at 4°C for 30min and bound with normalized virions at 4°C for 1 hour. Unbound virus was washed off and cell-bound virus was quantified using flow cytometry following sequential binding with rabbit anti-reovirus pAb and goat anti-rabbit Alexa Fluor 488. After secondary antibody staining, samples were fixed with 4% paraformaldehyde for 30 minutes at 4°C. Following a wash with PBS to remove 4% paraformaldehyde, cell pellet was gently resuspended in 500μl PBS. Samples were processed using a FACSCanto (BD Biosciences) and data was analyzed using FSC Express 5 (De Novo Software). Total cells were gated using FSC-A and SSC-A, while single cells were gated using FSC-A and FSC-H. A minimum of 10,000 total cells were collected for each sample. All steps were performed at 4°C on a microcentrifuge tube rotator and FACS buffer (PBS/5% FBS) was used as the diluent and wash buffer. Prior to fixation, cells were centrifuged at 500g for 5min at 4°C. After fixation, cells were centrifuged at 1,000g for 5min at 4°C. Two wash steps were performed following virus binding and primary and secondary antibody incubation. Note that a “wash” in the above protocol constitutes centrifugation, aspiration of supernatant, resuspension of pellet in wash buffer, centrifugation and aspiration of wash buffer.

### Flow cytometry cell death assay

Early apoptosis and cell death were measured by Annexin V and 7-AAD staining precisely as previously described (Shmulevitz and Lee, 2012), prior to processing with FACSCanto (BD Biosciences) and analysis with FSC Express 5 (De Novo Software). Identical gates were applied to all samples of a given experiment, and were applied based on discrimination of the live (Annexin V and 7-AAD negative) population.

### Agarose gel separation of reovirus

Purified virions (5×10^10^ virus particles) diluted in 5% Ficoll and 0.05% bromophenol blue were run on a 0.7% agarose gel in TAE buffer (40mM Tris, 5 mM sodium acetate, 1 mM EDTA [pH 7.5]) for 12 hours at room temperature, stained with Imperial total protein stain for 2hrs at room temperature, destained overnight in TAE buffer and visualized on the ImageQuant LAS4010 imager (GE Healthcare Life Sciences)

### In-vitro reovirus core transcription assay

Reovirus cores were generated by incubating purified virions with chymotrypsin (CHT) (C3142, Millipore Sigma) at 14μg/ml for 2 hours at 37C. CHT digest reactions were halted by adding protease inhibitor cocktail (11873580001, Roche) and incubating at 4°C. Reovirus cores were pelleted by centrifugation at 100,000g for 2 hours at 4°C, and reconstituted in 100mM Tris pH 8. Transcription reactions were assembled on ice to include 100mM Tris pH 8, 10mM MgCl2, 100μg/ml pyruvate kinase (P7768, Millipore Sigma), 3.3mM phosphoenol pyruvate (P0564, Millipore Sigma), 0.32 units/μl RNaseOUT (10777019, ThermoFisher Scientific), 0.2mM rATP, 0.2mM rCTP, 0.2mM rGTP, 0.2mM rUTP and 1×1011 virus cores per 150μl reaction. Negative control samples were set up without rATP. Reactions were allowed to proceed at 40°C and at indicated timepoints, 40μl transcription aliquots were added to 400ul TRI Reagent LS (T3934, Millipore Sigma) containing 3ng of mouse GAPDH RNA (in-vitro transcribed using T7 RiboMAX (Promega), as per manufacturer’s protocol). Using 10μg glycogen (R0551, ThermoFisher Scientific) as a carrier according to manufacturer’s instructions, RNA was purified, converted to cDNA (28025013, ThermoFisher Scientific) using random primers (48190011, ThermoFisher Scientific) and RT-PCR (1725204, Bio-Rad) performed to quantify reovirus S4, reovirus M2 and mouse GAPDH. Values were standardized to GAPDH and plotted relative to 0 hours post transcription. For high throughput transcription assays, reactions were set up similar, aliquoted into RT-PCR 96-well tubes and spiked with 10× final SYBR Green II (S7564, ThermoFisher Scientific). Relative fluorescence was measured at 5-minute intervals for 2 hours in a CFX96 system (Bio-Rad).

### In-vivo oncolysis experiments

Fifteen six-week-old female C57BL/6 mice (Charles River Laboratory, Montreal, Canada) were injected subcutaneously in the hind flank with 1×10^5^ B16-F10 cells per 100ul per mouse. When tumors become palpable (14 days post B16-F10 cell injection), a total of 3 equivalent doses (5×10^8^ pfu/100ul) were inoculated intratumorally at 2-day intervals. The negative control group was inoculated with PBS. Tumor volumes were measured in 3 dimensions using digital calipers every 2 days. Mice were sacrificed when either tumors became too large (200mm3), and/or tumors had visible signs of necrosis and ulceration. All experiments were done in accordance with guidelines on humane handling of experimental animals as established by the Canadian Council on Animal Care.

### T3D-PL and T3D-TD reverse genetics system

L929 cells in 6well plates were infected with T3D-PL or T3D-TD at an MOI of 3 and cell lysates were collected at 24hpi in TRI Reagent® (T9424, Millipore Sigma). Total RNA was purified using as per TRI Reagent® protocol. Using 1μg RNA per 20μl reaction, cDNA synthesis was performed with pooled forward and reverse gene specific primers for all 10 genes, and M-MLV reverse transcriptase (28025013, ThermoFisher Scientific) as per M-MLV reverse transcriptase protocol. Using cDNA (2μl) as a template, each gene segment was PCR amplified (100μl total reaction) using a gene specific primer set with iProof High Fidelity PCR kit (BioRad), as per manufacturer’s protocol. PCR products were purified using QIAquick PCR Purification Kit (28106, Qiagen). S1, M1, M2, M3, L1, L2 and L3 gene segments were gel extracted using PureLink™ Quick Gel Extraction Kit (K210012, Fisher Scientific). All purified PCR products were double digested with CpoI (FD0744, ThermoFisher Scientific) and NotI (FD0594, ThermoFisher Scientific) and purified using QIAquick PCR Purification Kit (28106, Qiagen).

Vector pBacT7-S1T3D (addgene plasmid #33282) was double digested with CpoI (FD0744, ThermoFisher Scientific) and NotI (FD0594, ThermoFisher Scientific), and vector backbone (5716bp fragment) was gel extracted using PureLink™ Quick Gel Extraction Kit (K210012, Fisher Scientific). Gel extracted vector backbone was treated using alkaline phosphatase (EF0654, ThermoFisher Scientific) as per manufacturer’s protocol, and purified using QIAquick PCR Purification Kit (28106, Qiagen).

Ligation reactions were performed with a 1:5 ratio of vector:insert using T4 DNA Ligase (15224017, ThermoFisher Scientific) in a 20ul reaction as per manufacturer’s protocol. Ligation reactions were transformed into Stbl3 chemically competent bacteria and transformants were selected on LB/agar/carbnecillin (100 μg/ml) plates. For each gene, 8 bacteria colonies were selected for colony PCR validation. Positive bacteria clones were inoculated into 50ml LB/carbnecillin (100 μg/ml) culture for amplification and midiprep plasmid purification using GenElute™ HP Plasmid Midiprep Kit (NA0200, Millipore Sigma). All plasmids except T3D-L1 were amplified at 37°C for 14-16hrs. T3D-L1 plasmids were amplified at 30°C for 24-28hrs. All plasmids were sequenced to validate inserted gene sequences.

### Reassortant reovirus generation

BHK-21-BSR T7/5 cells were seeded in 12 well plates and transfected when cells reached 80-90% confluency. Volumes below are for a single 12 well reaction: 0.35μg of each of 10 reovirus genes and 0.8μg C3P3 (T7 RNA Polymerase linked with (G4S)4 to NP8686 African Swine Fever Virus capping enzyme) were diluted in 50μl Opti-MEM (31985070, ThermoFisher Scientific). 11μl TransIT®-LT1 (MIR2300, Mirus Bio) was diluted in 50μl Opti-MEM and incubated for 10 minutes at room temperature. LT1 and plasmid dilutions were combined and incubated for 20 minutes at room temperature. Cell media was replaced with 500μl fresh media and transfection mixture was added dropwise to each well. Following a 16-18 hour incubation at 37°C, media was replaced with 750μl fresh media and incubated for an additional 3-5days. Lysates were scraped, collected and plaqued on L929 cells following three freeze/thaw cycles.

### Lentivirus production and generation of stable cell lines expressing shRIG-I, shEMPTY or shSCR for microarray and qRT-PCR experiments in supplemental figures

Lentivirus was generated as per MISSION® Lentiviral Packaging Mix protocol (SHP001, Millipore Sigma) using 6 well plates. Lentivirus containing supernatants were collected every 12hrs for 72hrs. Pooled lentivirus collections were centrifuged at 600g for 10min at 4°C, and the supernatant was 0.45μm filter sterilized, aliquoted and stored at −80°C.

Lentivirus stock was diluted in cell culture media supplemented with sequabrene (S2667, Millipore Sigma) at 8μg/ml final. Dilutions ranged from 1/3 to 1/36. Media was aspirated from 12 well plates with cells at 50-60% cell confluency, 500μl lentivirus dilution was added to each well and allowed to incubate at 37°C for 12hrs. Lentivirus was aspirated and replaced with cell culture media for an additional 12-24hrs until cells became confluent. At 100% confluency, cells were trypsinized, transferred to a 6 well plate and incubated at 37°C for 12hrs. Cell culture media was replaced with media supplemented with puromycin (2μg/ml for NIH/3T3 cell line) and cell death was monitored by microscopy. Cells not treated with lentivirus but exposed to puromycin supplemented media were used to determine when untransduced cells were killed. When lentivirus transduced cells reached 90-100% confluency, cells, were trypsinized and transferred to 55cm2 flask with media supplemented with puromycin. Lowest lentivirus dilution with minimal (0-10%) cell death was selected for further assessment. Puromycin selection was performed every second passage.

### Microarray Analysis

NIH/3T3 cells transduced with lentivirus generated from either empty plko.1 vector (shEMPTY) or plko.1 containing shRNA targeting mouse RIG-I (shRIG-I) were selected with 2μg/ml puromycin for seven days, followed by T3D^PL^ infection at the dose empirically determined to cause 70% reovirus-antigen-positive cells on NIH/3T3 shEMPTY cells by immunofluorescence with anti-reovirus polyclonal antibodies. At 12 hpi, RNA was extracted from reovirus infected and mock infected samples and sent for microarray processing at MOgene LC (1005 North Warson Road, Suite 403, St. Louis, MO 63132 USA) using the Agilent Mouse SurePrint G3 GE, 8×60k – 028005 array type. Raw microarray image files were extracted with Agilent’s Gene Expression Microarray Software, GeneSpring GX and is accessible from NCBI GEO DataSet # GSE35521. Data sets for untreated and LPS treated, WT, cRel−/− p65−/− and IFNAR−/− MEFs were obtained from NCBI GEO DataSet # GSE35521 (Cheng C.S. et al, 2011). For each data set, the normalized microarray data along with gene and sample information was input into the pheatmap package in R. Hierarchical clustering of data was performed with “Euclidean distance” as the measure of dissimilarity. Heatmaps were generated using pheatmap and the associated dendrogram clusters were extracted for further analysis.

## ACKNOWLEDGEMENTS

This work was funded by a Canadian Institutes of Health Research (CIHR) project grant MS, a project grant from the Li Ka Shing Institute of Virology to MS, a salary award to MS from the Canada Research Chairs (CRC) and infrastructure support from Canada Foundation for Innovation (CFI). AM received scholarships from the Alberta Cancer Foundation (ACF) and the Faculty of Medicine and Dentistry. We would like to thank Kevin Coombs at the University of Manitoba and Terrence Dermody at University of Pittsburgh for generously sharing their laboratory reovirus T3D virus lysates, Aja Reiger at the Faculty of Medicine and Dentistry flow cytometry facility, Rob Maranchuk at the Li Ka Shing Institute of Virology RNAi screening facility and Stephen Ogg at the Faculty of Medicine & Dentistry cell imaging centre, for valuable technical advice, training and support. These core facilities at the University of Alberta receive financial support from the Faculty of Medicine and Dentistry and Canadian Foundation for Innovation (CFI) awards. We appreciate all the helpful discussions and suggestions by members of the Maya Shmulevitz, David Evans, Mary Hitt, Ronald Moore and Patrick Lee laboratories.

## AUTHOR CONTRIBUTIONS

Conceptualization, A.M. and M.S.; Methodology, A.M and M.S.; Investigation, A.M., D.R.C., P.K. and M.S.; Writing – Original Draft, A.M. and M.S.; Writing -Review & Editing, A.M., J.R.S. and M.S.; Funding Acquisition, M.S.; Resources, M.E.V and C.K.B.; Supervision, M.S., J.R.S., S.A.G.

## SUPPLEMENTAL INFORMATION TITLES AND LEGENDS

**Figure S1: Virus-cell binding quantification using flow cytometry. (A)** L929 cells were detached non-enzymatically using cell dissociation buffer (Sigma, C5914). T3D^PL^ dilutions were bound at 4°C, and following extensive washing to remove unbound reovirus, cell-bound reovirus was stained using polyclonal reovirus antibodies and mean fluorescence intensity (MFI) quantified using flow cytometry (BD FACSCanto). Black histogram = MOCK, Dark orange = highest reovirus dilution, Light orange = lowest reovirus dilution. **(B)** Linear regression analysis of MFI determined from (A), corrected for mock infected background (n = 2).

**Figure S2: T3D^PL^ establishes a productive infection earlier than T3D^TD^**. **(A)** L929 cells were mock infected or infected with T3D^PL^ or T3D^TD^ at the indicated MOI (based on titers established on L929 cells) and incubated at 37°C for 12, 24 or 48 hours. Paraformaldehyde-fixed cells were then subjected to immunofluorescence staining with reovirus specific primary antibody and Alexa Fluor 488 conjugated secondary antibody as described in materials and methods of the main manuscript. Nuclei were stained with HOESCHT 33342. Immunofluorescence was imaged on the EVOS FL Auto Cell Imaging System. **(B)** Similar to (A) except at each time point, cells were subjected to immunofluorescence staining and flow cytometric analysis as described in materials and methods of the main manuscript. Gates were set based on Mock-infected cells. **(C)** Similar to (B) but on H1299, ID8, and B16-F10 cancer cell lines.

**Figure S3: RIG-I signaling does not impact initial reovirus infection.** NIH/3T3 cells stably transduced with scrambled (shSCR) or RIG-I (shRIG) lentivirus were mock-infected or infected with reovirus at MOIs of 20 and 60 (relative to titers obtained on the more-susceptible L929 cells), and incubated at 37°C. **(A)** At 12hpi, RNA from cells infected at MOI of 20 was extracted and subjected to cDNA synthesis and qRT-PCR for mouse RIG-I (Rig-i) and mouse GAPDH. RIG-I mRNA levels were corrected for GAPDH and presented relative to shSCR T3D^TD^ (set to 1.0). Two independent experiments are each represented by individual dots. **(B)** At 12hpi, samples were subjected to Western Blot analysis for RIG-I, reovirus μ1 and σ3, or β acting loading control. **(C)** At 12 and 24hpi, cells underwent immunofluorescence staining with reovirus specific primary antibody and Alexa Fluor 488 conjugated secondary antibody to detect reovirus infected cells (Green), and HOESCHT 33342 to detect nuclei (Blue). Green staining represents reovirus infected cells, and blue staining represents cell nuclei. **(D)** Similar to (A) but qRT-PCR was conducted for viral and cellular genes indicated above each plot, and following infection at MOIs of both 20 and 60 as indicated.

**Figure S4: Characterization of RIG-I and NFκB dependent reovirus induced genes.** A description of the microarray data used in this figure is provided in supplemental materials and methods. A list of candidate genes was selected based on the following criteria: (1) Genes induced (≥ 2-fold) following T3D^PL^ infection in shEMPTY NIH/3T3 cells, and (2) Genes inhibited (≥ 2-fold) in cRel -/- p65-/- MEFs treated with LPS, relative to WT MEFs treated with LPS. **(A)** Hierarchical clustering heat maps depicting gene subsets dependent on RIG-I, IFNAR and/or NFκB as described below for each cluster. **(B)** Collection of graphs showing gene clusters from (A). Cluster 1: reovirus induced genes dependent on RIG-I signaling, based on ≥ 2-fold change between shEMPTY versus shRIG-I. Cluster 2: reovirus induced genes independent of RIG-I signaling, based on, based on < 2-fold change between shEMPTY versus shRIG-I. Cluster 3: reovirus induced genes dependent on RIG-I and NFκB signaling but independent of IFNAR signaling, based on genes in Cluster 1 that also change ≥ 2-fold between WT versus cREL^−/-^p65^−/-^ but < 2-fold change between WT and IFNAR^−/-^. Cluster 4: reovirus induced genes dependent on RIG-I, IFNAR and NFκB signaling, based on genes in Cluster 1 that also change ≥ 2-fold between WT and both cREL^−/-^p65^−/-^ and IFNAR^−/-^. Cluster 5: reovirus induced genes independent of RIG-I signaling but dependent on IFNAR and NFκB signaling, based on genes in Cluster 2 that also change ≥ 2-fold between WT and both cREL^−/-^p65^−/-^ and IFNAR^−/-^. Cluster 6: reovirus induced genes independent of RIG-I and IFNAR signaling but dependent on NFκB signaling, based on genes in Cluster 2 that also change ≥ 2-fold between WT versus cREL^−/-^ p65^−/-^ but < 2-fold change between WT and IFNAR^−/-^.

**Figure S5: Classification of IFN-dependent and IFN-independent genes and NFκB modulation effects on reovirus**. **(A)** L929 cells were treated with 1/10 dilutions (initial 1000 U/ml/12well) of purified IFNα or IFNβ for 12 hours at 37°C. Samples were collected for RNA extraction, cDNA synthesis and RT-PCR using gene-specific primers (corrected for GAPDH). Values were standardized to untreated sample, n = 3. **(B, Left)** L929 cells were either mock infected or infected with T3D^PL^ or T3D^TD^ at an MOI of 3 and co-incubated throughout infection with either InSolution™ NF-κB Activation Inhibitor (Calbiochem, 481407) at 10nM, 2nM or 0.4nM, mouse TNF (Sigma T7539) at 10ng/ml, 2ng/ml, or 0.4ng/ml, or left untreated (UT). At 24hpi, cells were stained for cell death using Annexin V and 7-AAD as described in materials and methods in the main manuscript. Cells were considered ‘dying cells’ if positive for either Annexin V and 7-AAD relative to untreated and mock infected cells. N=3, 2way ANOVA with Dunnett's multiple comparisons test, ***p<0.001. **(B, Right)** Similar to (B, Left), however cells were either pre-treated for 3 hours with 10nM InSolution™ NF-κB Activation Inhibitor prior to infection and co-infection treatments, or not treated prior to infection. Co-infection treatments included InSolution™ NF-κB Activation Inhibitor at 10nM, 2nM, 0.4nM or 0.08nM, mouse TNF at 10ng/ml, 2ng/ml, 0.4ng/ml, or 0.08ng/ml, or left untreated (UT) **(C)** Similar to (B, Right) except that samples were collected at 12hpi for Western Blot analysis to detect levels of reovirus proteins (μ1 and σ3), IKKα, NFκB-induced gene JunB, or β-Actin loading control, for InSolution™ NF-κB Activation Inhibitor at 10nM, 2nM, and 0.4nM or mouse TNF at 10ng/ml and 2ng/ml.

**Figure S6: Crystal structures highlighting T3D^PL^ and T3D^TD^ polymorphisms in S4 and L3 genes**. All structures were visualized and modified using the PyMOL molecular visualization program by Schrodinger. **(A)** σ3 dimer (1fn9.pdb). Ribbons in shades of blue indicate 2 monomers of σ3. Polymorphisms in σ3 are represented by yellow spheres (W133R), red spheres (G198K), pink (E229D). Regions previously determined to bind dsRNA are indicated in cyan spheres. **(B)** σ33μ13 heterohexamer (1jmu.pdb). Ribbons in shades of blue indicate 3 monomers of σ3 and shades of green indicate 3 monomers of μ1. Polymorphisms in σ3 are represented by yellow spheres (W133R), red spheres (G198K), pink (E229D) and orange spheres (Y354H).

**Table S1: Key Resources Table**

**Table S2: Accession numbers for sequences analyzed in the manuscript**

**Table S3: Oligonucleotide sequences for cloning reovirus genes into the reverse genetics backbone plasmid**

**Table S4: Oligonucleotide sequences for primers used in qRT-PCR**

**Table S5: Oligonucleotide sequences for primers used for site-directed mutagenesis**

**Table S6: Lists for all genes in clusters 1-6 described in Figure S4**

